# Hydroxytyrosol (HT) analogues act as potent antifungals by direct disruption of the fungal cell membrane

**DOI:** 10.1101/350025

**Authors:** George Diallinas, Nausica Rafailidou, Ioanna Kalpaktsi, Aikaterini Christina Komianou, Vivian Tsouvali, Iliana Zantza, Emmnauel Mikros, Alexios Leandros Skaltsounis, Ioannis K. Kostakis

## Abstract

Fungal infections constitute an emerging threat and a prevalent health problem due to increasing number of immunocompromised people and pharmacological or other treatments aiming at viral infections, cancer or allergies. Currently used antifungals suffer from inefficiency, toxic side effects and developing drug-resistance. Additionally, over the last two decades no new classes of antifungals have been approved, emphasizing the urgent need for developing a novel generation of antifungals. Here we investigate the antifungal activity of a series of chemically synthesized Hydroxytyrosol (HT) analogues. HT is one of the major phenolic compounds in olive oil, shown to possess radical-scavenging antioxidant, antiproliferative, proapoptotic and anti-inflammatory activities. No previous report has studied HT analogues as antifungals. We show that specific analogues have broad and strong antifungal activity, significantly stronger than the parent compound HT. Using *A. nidulans* as an *in vivo* cellular model system, we show that antifungal HT analogues have an unprecedented efficiency in fungal plasma membrane destruction. Importantly, antifungal HT analogues did not show toxicity in a mammalian cell line, whereas no resistance to HT analogues was obtained by standard mutagenesis. Our results open the way for the development of a novel, efficient and safer class of antifungals.

## Introduction

In recent years, systemic fungal infections have emerged as an increasingly prevalent health problem (McCarthy et al., 2017). Infections are rising among immunocompromised patients, including individuals suffering from HIV/AIDS or diabetes mellitus, or following organ transplantations and immunosuppressive chemotherapy during cancer treatment (Low and Rotstein 2011). The most clinically significant invasive opportunistic fungal pathogens belong to one of the four groups: *Aspergillus, Candida, Cryptococcus* and *Pneumocystis*, with the first two being the most important of all fungal pathogens (Köhler et al., 2017). Currently used antifungals include three major classes of drugs with different mechanisms of action: polyenes (disrupt fungal membranes), azoles (inhibit ergosterol biosynthesis), and echinocandins (inhibit synthesis of cell wall β-glucan) (Odds et al., 2003). However, all current antifungals suffer from inefficiency, toxic side effects, drug-drug interactions and developing drug-resistance (Arendrup 2014; Cuenca-Estrella 2014; Fairlamb et al., 2016). In addition, since 2001, no new classes of antifungals have been approved. This emphasizes the urgent and critical need for developing a novel generation of antifungals.

Pharmacological therapies for various bacterial or viral infections based on natural, mostly herbal, agents are among the most current therapeutic trends in medicine (Pan et al., 2013; Li and Weng 2017). Among these, a selection of Mediterranean plant extracts from olive (*Olea europaea*), parsley (*Petroselinum crispum)*, oregano (*Origanum vulgare*), thyme (*Thymus vulgaris*), sage (*Salvia officinalis* and others have shown a significant inhibitory activity against numerous bacteria, while antiviral lectins from red and blue-green algae show potent *in vitro* and *in vivo* activity against hepatitis C virus (Bower et al., 2016). Recently, plant extracts and chemically synthesized related analogues from olive have also shown an antiprotozoan activity (Belmonte-Reche et al., 2016; Koolen et al., 2016). Hydroxytyrosol (HT), one of the major phenolic compounds in olive oil, has recently received particular attention because of its radical-scavenging, antiproliferative, proapoptotic and anti-inflammatory activities, which seem to have a counteractive effect on carcinogenesis and other cellular malfunctions in animal trials and *in vitro*. Additionally, recent evidence has shown that HT and its analogues might be promising antibacterial, antiviral or antiprotozoan agents (Manna et al., 2005; Fernandez-Bolanos et al., 2008; 2012; Chillemi et al., 2010; Koolen et al., 2016; Thielmann et al.,2017; Robles-Almazan et al., 2018). However, no systematic effort has been invested in search of novel antifungals based on HT, except from some reports concerning yeast species (Pereira et al., 2007; Zoric et al., 2013), or HT analogues.

Based on some preliminary tests of our lab that showed a moderate antifungal effect of HT and some HT analogues on *Aspergillus nidulans*, here we systematically synthesize and test the antifungal activities of an extended series of novel HT analogues. The rationale of their synthesis was based mostly on the possible effect of the length and saturation of the fatty acid chain, and the substitution of the α-carbon of the HT side chain. We show that several of the synthesized HT analogues have a very potent and broad antifungal action against major filamentous fungal pathogens (*Aspergillus fumigatus, Aspergillus flavus, Fusarium oxysporum*), *Aspergillus nidulans* and *Candida albicans*. Importantly, we reveal that the antifungal activity of HT analogues is directly related to its rapid destructive effect on fungal plasma membranes, which in turn justifies why resistance mutants could not be isolated. Given that we also obtained preliminary evidence that HT analogues do not show cytotoxicity against a mammalian cells line, our work is expected to open the way for the developing of a new class of highly potent novel antifungals.

## Results

### Chemical synthesis of HT analogues

23 analogues of HT were synthesized as described in detail in Materials and methods and in Supplementary material (**Supplementary Figure 1**). The new compounds are lipophilic esters of HT bearing modifications on the α-carbon of the catechol side chain. More precisely, the new compounds are categorized in 3 different series (**Figure 1a**). The compounds of the second series are simple esters of HT in the aliphatic hydroxyl group, whereas the compounds of the fisrt series possess a carbonyl group on the α-carbon of the catechol side chain. The compounds of the third series possess a hydroxyl group on the α-carbon of the catechol side chain. In brief, chloride **1** was treated with the sodium salt of the appropriate acid to afford the required keto esters **2-12** (**Figure 1b**). The majority of the acids are commercially available, though the acids for the preparation of ester **9** was synthesized through a Wittig reaction of methyl 4-methylenecyclohexanecarboxylate. Treatment of the previous compounds with triethylsilane in trifluoroacetic acid, provided the desired, fully reduced, lipophilic esters **13-20**. It is noteworthy that this method was successful in yields up to 80%, with high reproducibility and scale up to grams. Finally, partial reduction with hydrogenation over Pd/C, provided the hydroxyl derivatives **21-24**, as racemic mixtures.

**Figure 1.**
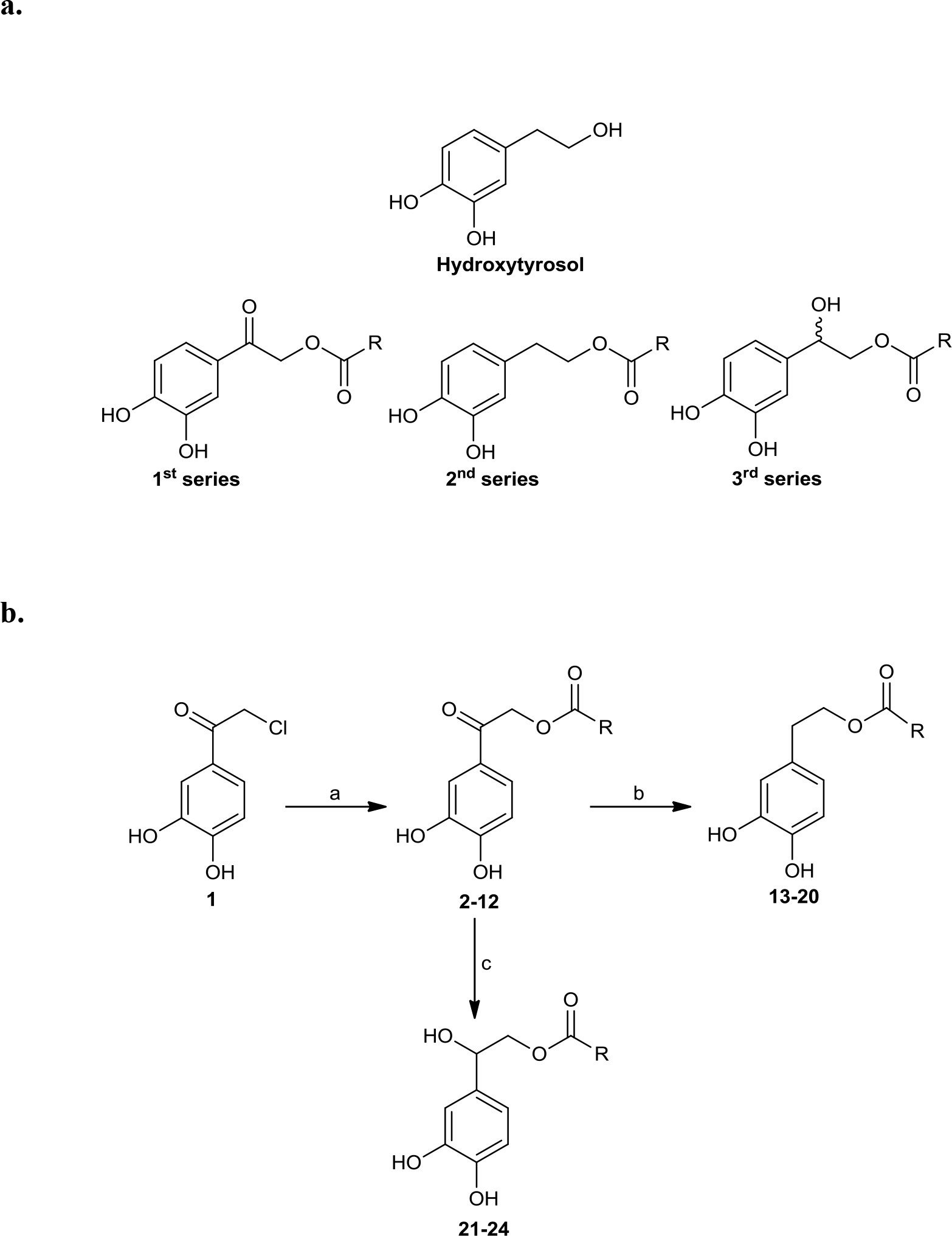
**a.** Structures of Hydroxytyrosol (HT) and general formula of the compounds **b.** Reactions and Conditions: a) sodium salt of suitable acid, DMF, 70 °C; b) Et3SiH, CF3COOH, r.t.; c) H2, 10% Pd/C, t-BuOH, 50 psi, r.t.

### HT analogues show high potential as antifungals

The 23 chemically synthesized analogues of HT were tested for their antifungal potential against *A. nidulans* and major fungal pathogens of animals (*A. fumigatus, C. albicans*) or plants (*A. flavus, F. oxysporum*). The rationale for choosing *A. nidulans* was based on its unique amenability as a model system for genetic and functional analysis, rather than its pathogenic profile, which would allow the investigation of the molecular mechanisms underlying of the antifungal action of HT analogues.

All synthesized HT analogues were tested on solid minimal media (MM) plus necessary supplements at physiological and optimal pH ranges (5.5-6.8) and temperatures (25-37 °C), for each of the five fungi chosen as targets. Growth rates and colony morphology was recorded after 2, 4 and 6 days. Initially, HT analogues were tested at a final concentration of 100 or 400 μM. The effect on *C. albicans* was recorded in liquid fresh cultures at their logarithmic phase of growth, but also on solid agar plates (for more details see Materials and methods). Similar results were obtained at the different pHs tested and on complete media or minimal media. Noticeably, recorded apparent antifungal activities were significantly higher at 37 °C compared to 25 °C (see later).

**Figure 2** highlights our findings and reflects the outcome obtained through several growth tests. Nine HT analogues (**2**, **4**, **5**, **10**, **11**, **15**, **16**, **18** and **19**), shown in Figure 3 had strong antifungal activity against *A. nidulans* at 37 ° C, mostly evident at 400 μM, and six of them (**2**, **5**, **11**, **15**, **16** and **19**) were also very active at 25 °C. Most of the same nine analogues also had strong antifungal activity against other fungi tested, at 25 °C and 37 °C (**Figure 2a** and test not shown). In particular, all nine analogues were extremely toxic to *A. fumigatus*, leading to total or extremely severe inhibition of growth at 100 μM. *A. flavus* proved to be the most resistant fungus among those tested, but still several analogues were highly inhibitory for its growth (**2**, **5**, **11**, **15**, **16** and **19**). Best antifungal agents against *F. oxysporum* proved to be analogues **2**, **5**, **11** and **15**. All analogues, at 100 μM, severely inhibited *C. albicans* growth in liquid cultures (**Figure 2b**) or on solid minimal media (not shown) at 37 °C. When liquid cultures were left to grow for more than 24 hours after the initial addition of HT analogues, growth *C. albicans* resumed in several cases, but not in the presence of analogues **4** or **15** (**Supplementary Figure 2**). This indicates that these analogues had the strongest cytotoxic effect or that these compounds were the most stable under the relative experimental conditions.

**Figure 2.**
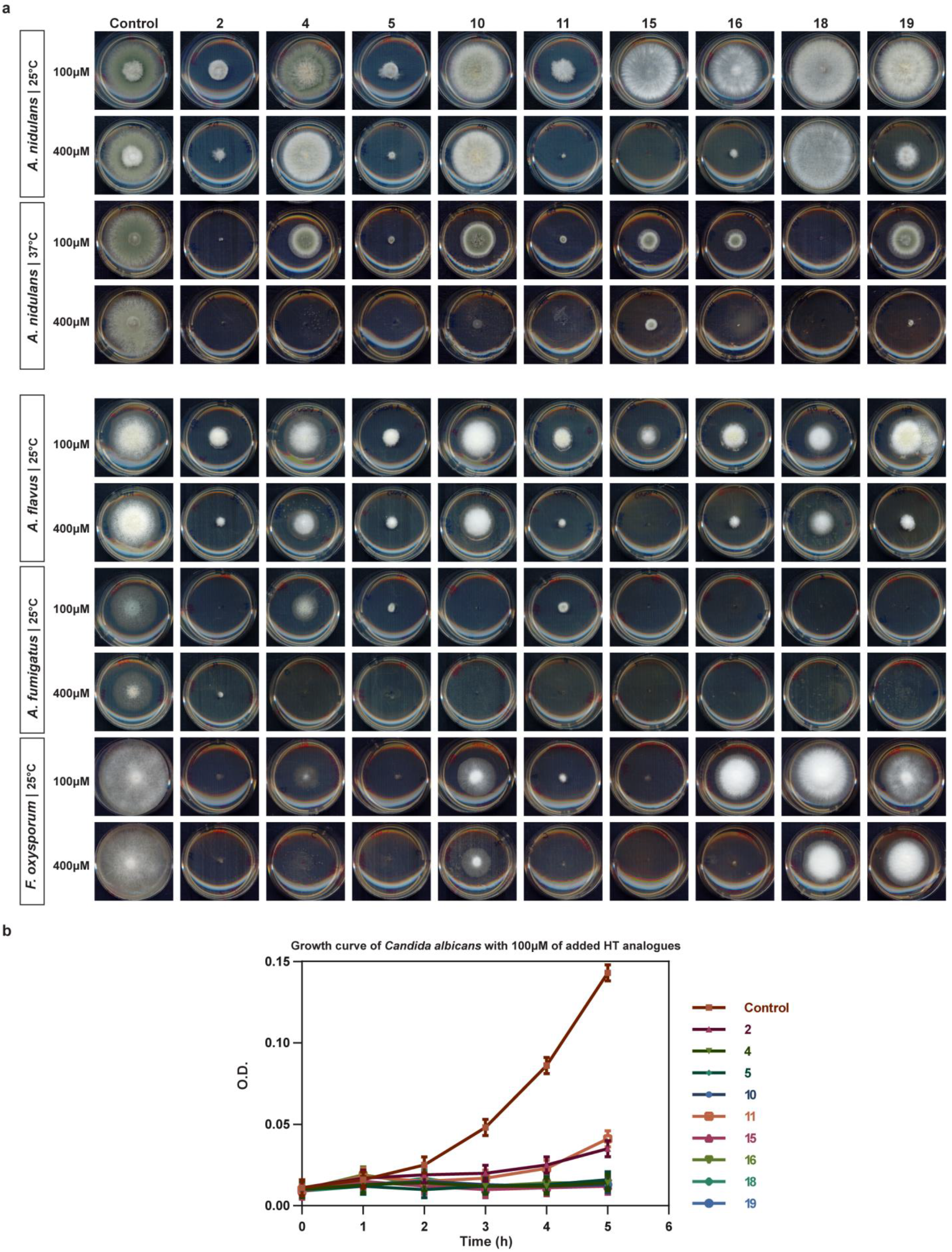
In vivo evaluation of HT analogues as antifungals. **a.** Growth tests showing the antifungal activity of certain HT analogues against pathogens *A. flavus A. fumigatus, F. oxysporum* and *A. nidulans*. Growth on two concentrations of HT analogues for each microorganism is shown. **b.** Growth curve of *C. albicans*. O.D stands of Optical Density at 600 nm of liquid cultures recorded hourly. Analogues were added to the cultures at 200 μM. Control stands for cultures were only DMSO solvent was added in the cultures, at the same concentrations as the analogues.

**Figure 3.**
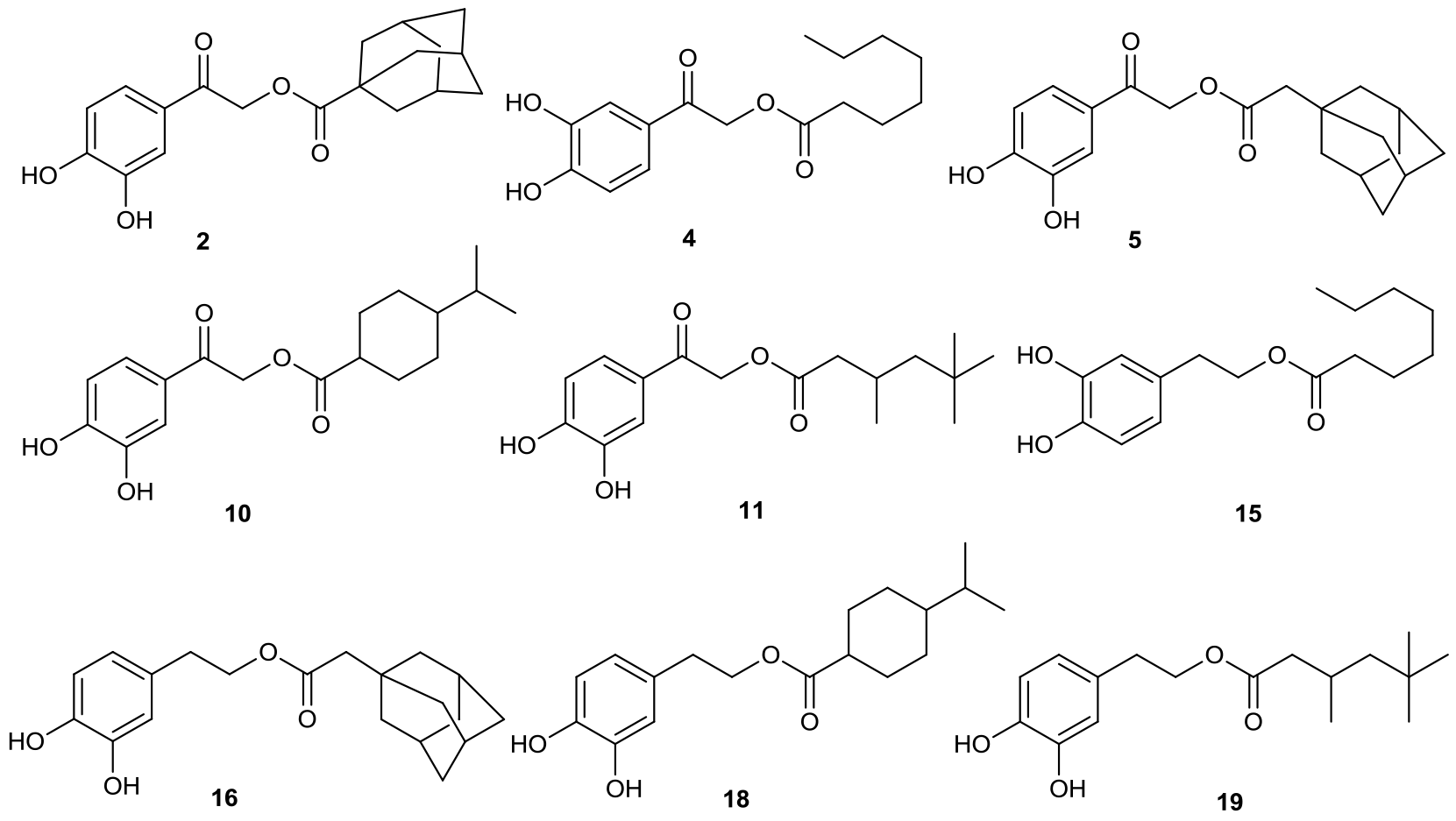
Specific HT analogues showing strongest antifungal activity. To further define the potential of HT analogues as antifungals, we tested their antifungal activity at a range of concentrations from 50-1000 μM. **Table 1** summarizes the results obtained, reflected as approximate MICs, ranging from <100 to <1000 μM, and IC_50_ values in the range 50-200 μM. Overall, **analogues 2**, **5**, **11**, **15**, **16** and **19** showed the most promising antifungal activity against all fungi tested. Notably, all of the selected analogues tested in **Figure 2** and **Table 1** possessed broad and strong antifungal activity, significantly higher than that of the parental natural product HT. Additionally, HT analogues had apparently stronger (i.e. against *A. nidulans*) or similar (against pathogenic fungi) antifungal activity compared to amphotericin B, a currently approved and administrated drug that acts on fungal membranes (see **Supplementary Figure 3**). The significance of the latter comparison will become apparent later.

**Table 1.**
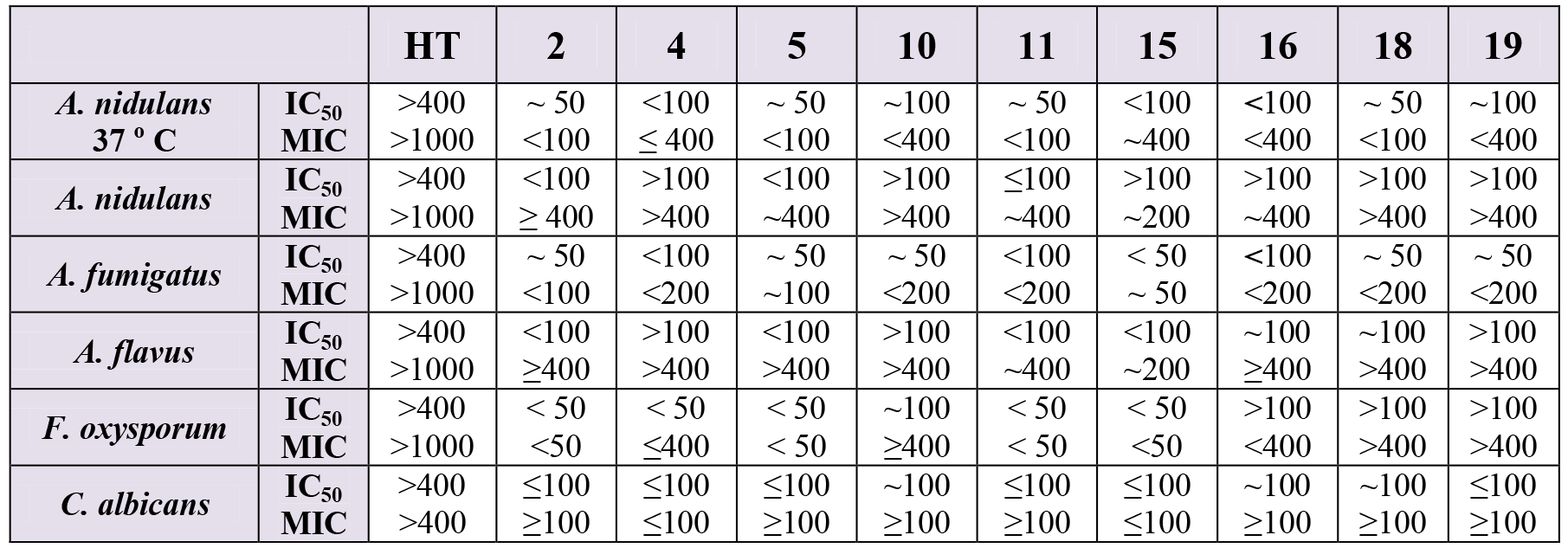
*MIC* and *IC*_50_ values of HT analogues showing antifungal activity. IC_50_ values shown correspond to the approximate μM concentration of the compounds that reduce the diameter of colony growth by 50% after 4-6 days at 37 ° C, on standard agar minimal media. MIC is the approximate μM concentration of HT analogues that lead to total inhibition of fungal vegetative growth under the same condition IC_50_ values were established. IC_50_ and MIC values are estimated by testing growth in the presence of a range of μM concentration (i.e. 0, 50, 100, 200, 400, 500 and 1000 μM). The values shown were estimated by at least three independent experiments, which showed no significant variation. For one of the most active analogues, **15**, additional experiments performed at a lower concentration range (5, 10, 20, 50 and 100 μM), led to a more precise estimation of IC_50_, which corresponds to 17 μM.

|  |  | HT | 2 | 4 | 5 | 10 | 11 | 15 | 16 | 18 | 19 |
| --- | --- | --- | --- | --- | --- | --- | --- | --- | --- | --- | --- |
| *A. nidulans*<br>37 ° C | IC<sub>50</sub> | >400 | ~ 50 | <100 | ~ 50 | ~100 | ~ 50 | <100 | <100 | ~ 50 | ~100 |
|  | MIC | >1000 | <100 | ≤ 400 | <100 | <400 | <100 | ~400 | <400 | <100 | <400 |
| *A. nidulans* | IC<sub>50</sub> | >400 | <100 | >100 | <100 | >100 | ≤100 | >100 | >100 | >100 | >100 |
|  | MIC | >1000 | ≥ 400 | >400 | ~400 | >400 | ~400 | ~200 | ~400 | >400 | >400 |
| *A. fumigatus* | IC<sub>50</sub> | >400 | ~ 50 | <100 | ~ 50 | ~ 50 | <100 | < 50 | <100 | ~ 50 | ~ 50 |
|  | MIC | >1000 | <100 | <200 | ~100 | <200 | <200 | ~ 50 | <200 | <200 | <200 |
| *A. flavus* | IC<sub>50</sub> | >400 | <100 | >100 | <100 | >100 | <100 | <100 | ~100 | ~100 | >100 |
|  | MIC | >1000 | ≥400 | >400 | >400 | >400 | ~400 | ~200 | ≥400 | >400 | >400 |
| *F. oxysporum* | IC<sub>50</sub> | >400 | < 50 | < 50 | < 50 | ~100 | < 50 | < 50 | >100 | >100 | >100 |
|  | MIC | >1000 | <50 | ≤400 | < 50 | ≥400 | < 50 | <50 | <400 | >400 | >400 |
| *C. albicans* | IC<sub>50</sub> | >400 | ≤100 | ≤100 | ≤100 | ~100 | ≤100 | ≤100 | ~100 | ~100 | ≤100 |
|  | MIC | >400 | ≥100 | ≤100 | ≥100 | ≥100 | ≥100 | ≤100 | ≥100 | ≥100 | ≥100 |

### Antifungal HT analogues show variable antibacterial activities

The HT analogues with the highest antifungal activity were also tested, at a range of 100-200 μM, in fresh exponentially growing bacterial cultures, for their potential as antibacterial agents. **Figure 4** shows the results obtained with *E. coli* and *B. subtilis*, as typical representatives of G- and G+ bacteria. All analogues were highly toxic to *B. subtilis* at 100 μM, but not at all to *E. coli. Pseudomonas* species showed differential growth behavior, with *P. aeruginosa* being fully resistant, but *P. fluorescens* highly sensitive at 200 μM (**Supplementary Figure 4**). Additional bacterial species were also tested (*Klebsiella, Enterococcus Staphylococcus, Streptococcus* and other *G-enterobacteria*) showing varying degrees of sensitivity to HT analogues, but this will be reported elsewhere as the present work is directed towards the discovery of novel antifungals.

**Figure 4.**
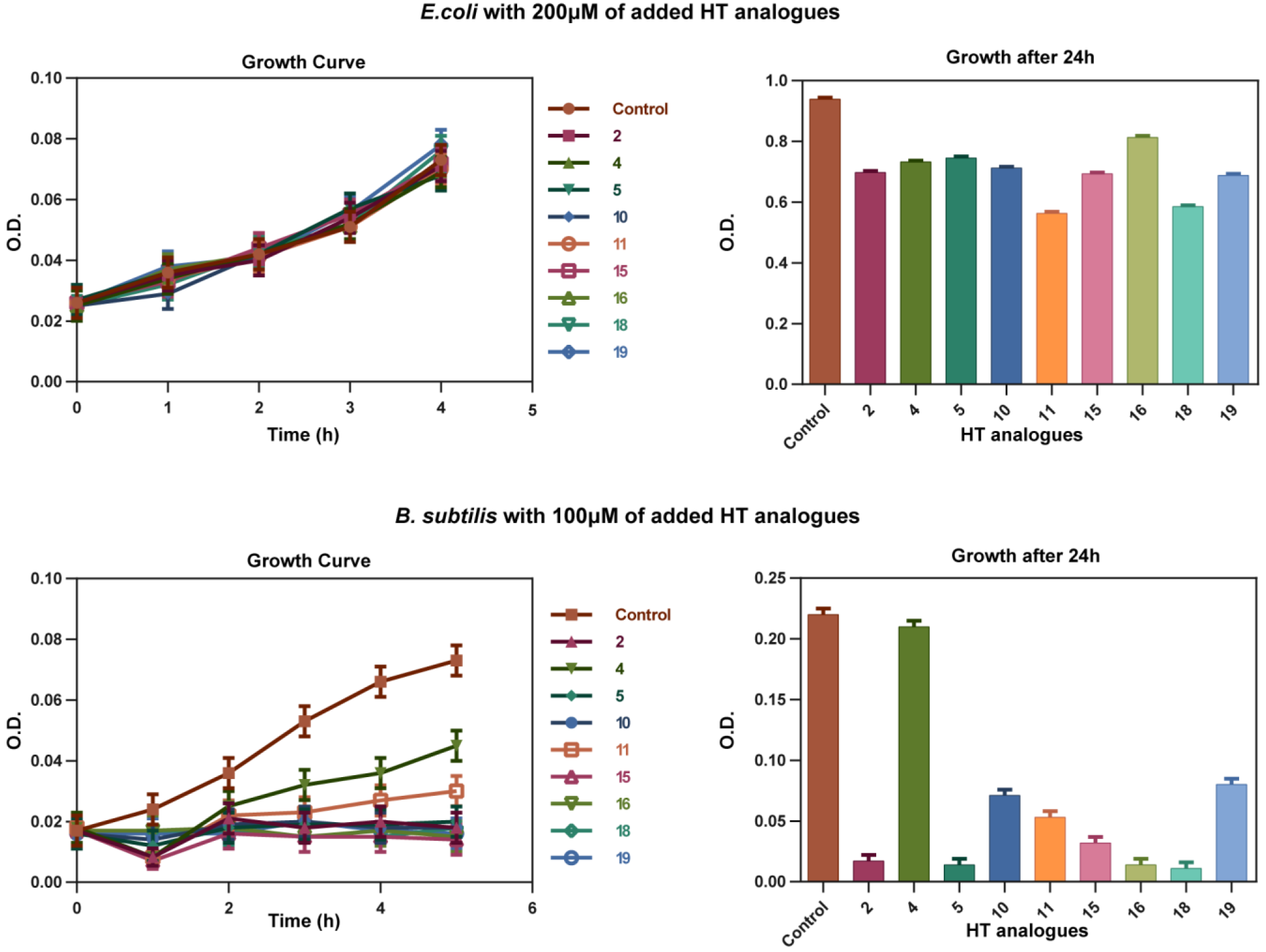
Antibacterial action of HT analogues against *E. coli* and *B. subtilis*. Growth curves on the left show O.D. values recorded hourly at 600 nm. Column bar graphs on the right show growth after 24 hours after the HT analogues addition, at 600 nm. Control stands for samples where only the solvent DMSO was added at a concentration identical to the one used for the dissolved analogues.

### Antifungal HT analogues disrupt the structure and function of *A. nidulans* plasma membrane

The non-dependence of the antifungal action of the HT analogues on whether minimal or complete media are used, or the pH range, when this was kept within limits proper for fungal growth, suggested that these compounds might be either taken up by nonfacilitated diffusion, or exert their activity directly without the need to enter the cell (i.e. on the cell wall or the plasma membrane). The relatively increased activity observed at 37 versus 25 ° C in the case of *A. nidulans* (see **Figure 2**) does not distinguish between the two possibilities, as increased membrane fluidity at a higher temperature would favor diffusion, as well as, binding of hydrophobic HT analogues in specific lipids of the fungal membrane. Thus, to investigate this issue directly we followed the effect of all antifungal HT analogues on the microscopic morphology and the plasma membrane (PM) of *A. nidulans* hyphae, using Brightfield and Epifluorescence microscopy, respectively. For investigating the effect on the plasma membrane we used strains expressing two GFP-tagged plasma membrane transporters, namely the UapA uric acid-xanthine transporter (Pantazopoulou et al., 2007) or the FurA allantoin transporter (Krypotou et al., 2015). **Figure 5** summarizes our results. Most HT analogues, when added at final concentration as low as 37.5 μM, for 0-30 min, had a rapid and prominent effect on plasma membrane integrity, reflected in dramatic reduction of transporter-associated peripheral GFP fluorescence signal and concomitant appearance of static, non-cortical membrane fluorescent aggregates (most evident with analogues **2**, **5**, **10**, **11**, **16** or **18**). Under brightfield we did not notice any dramatic modification that would be compatible with disruption of the cell wall or overall hyphae morphology (see also **Supplementary Figure 5**), despite a visible increase in vacuole number and size. These effects were practically immediate, becoming evident in 1-5 min, which also somehow excludes the idea that HT analogues act primarily by metabolic inhibition of an enzyme. In general, the relative strength of the detrimental effect of the different HT analogues on the PM was variable, with some analogues leading to total apparent disintegration, while others led to significant but not total, of the PM in 10 min. The concentration used to test the analogues was < 50 μM (37.5 μM) in order to avoid any effect of DMSO (the solvent used) on the stable localization of transporters in the PM, as we have noticed that DMSO concentrations > 50 μM elicit a degree of endocytic turnover for most transporters studied in our lab (unpublished observations).

**Figure 5.**
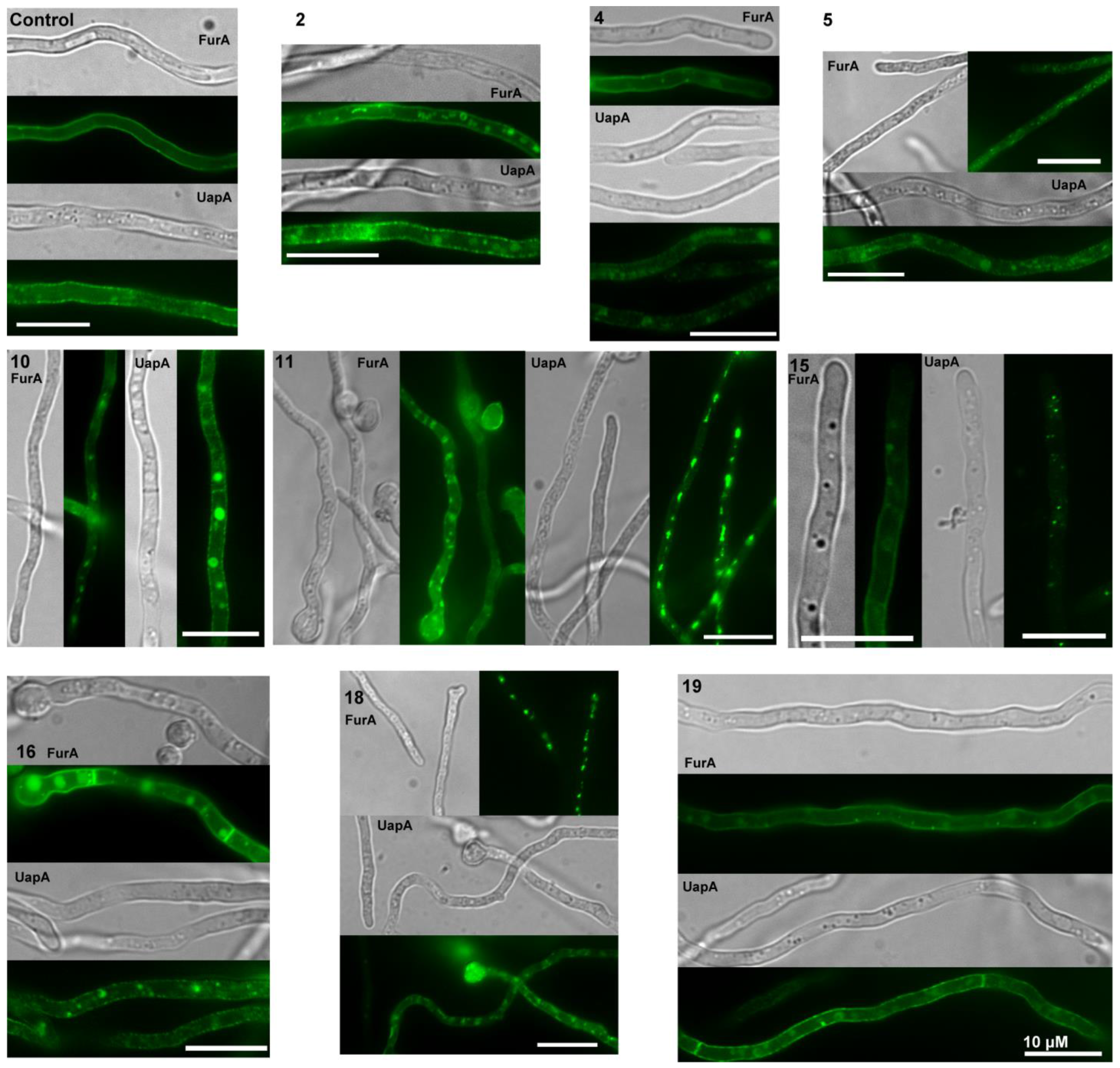
Effect of HT analogues on *A. nidulans* plasma membrane and transporter-mediated uptake of xanthine. Epifluorescence *in vivo* microscopy showing the effect of HT analogues (37.5 μM) on the plasma membrane of *A. nidulans*. The picture shows young hyphae of strains expressing functional GFP-tagged FurA or UapA transporters as PM molecular markers. FurA expression is stronger than that of UapA, due to transcription via a strong constitutive promoter (*gpdA*_*p*_), while UapA is transcribed by its native, relatively weak, promoter. Notice that upon addition of most analogues cortical fluorescent labeling is reduced with concurrent appearance of cytosolic fluorescent foci, which represent membrane aggregates. Scaleσ shown are 10 μM.

To further examine the nature of the effect of HT analogues on the PM, we also performed direct transport assays of radiolabeled metabolites, imported by specific transporters, in the presence of excess analogues. Transport assays used measure initial uptake rates (at 60 sec) of radiolabelled substrates in germinated conidiospores (Krypotou and Diallinas, 2014). In particular, we tested the uptake of radiolabeled xanthine, which is specifically transported by two uptake systems, the UapA (~70%) and UapC (~30%) transporters (Pantazopoulou and Diallinas 2007; Krypotou and Diallinas, 2014). Results are summarized in **Figure 6**. HT analogues **2**, **4**, **5**, **10**, **11**, **15** and **16** reduced xanthine uptake to ~20-40% of the control sample (only DMSO added), whereas the rest (**18** and **19**) showed less inhibitory effect. Overall, these results strongly suggested that most HT analogues tested had a rapid negative effect on fungal transport systems. Given that the relevant xanthine transporters, UapA and UapC, as most fungal transporters, are H^+^ symporters, our results can in principle be explained by two scenarios; either the analogues lead to rapid disorganization of the PM, or they led to a rapid depolarization of the membrane, acting as direct H^+^ gradient uncouplers. However, the latter case seems unlikely, or secondary, under the light of the microscopic analysis shown in **Figure 5**, which directly confirmed the dramatic effect of all HT analogues tested on the PM within some minutes after their addition to the fungal cultures. Additionally, the strength of inhibition of transporter-mediated xanthine uptake by the different HT analogues tested was in good agreement with the results obtained following their effect on PM integrity (**Figure 5**) and their *in vivo* antifungal activity (**Figure 2**). We thus conclude that the direct target of HT analogues is disruption of the fungal PM integrity and function.

**Figure 6.**
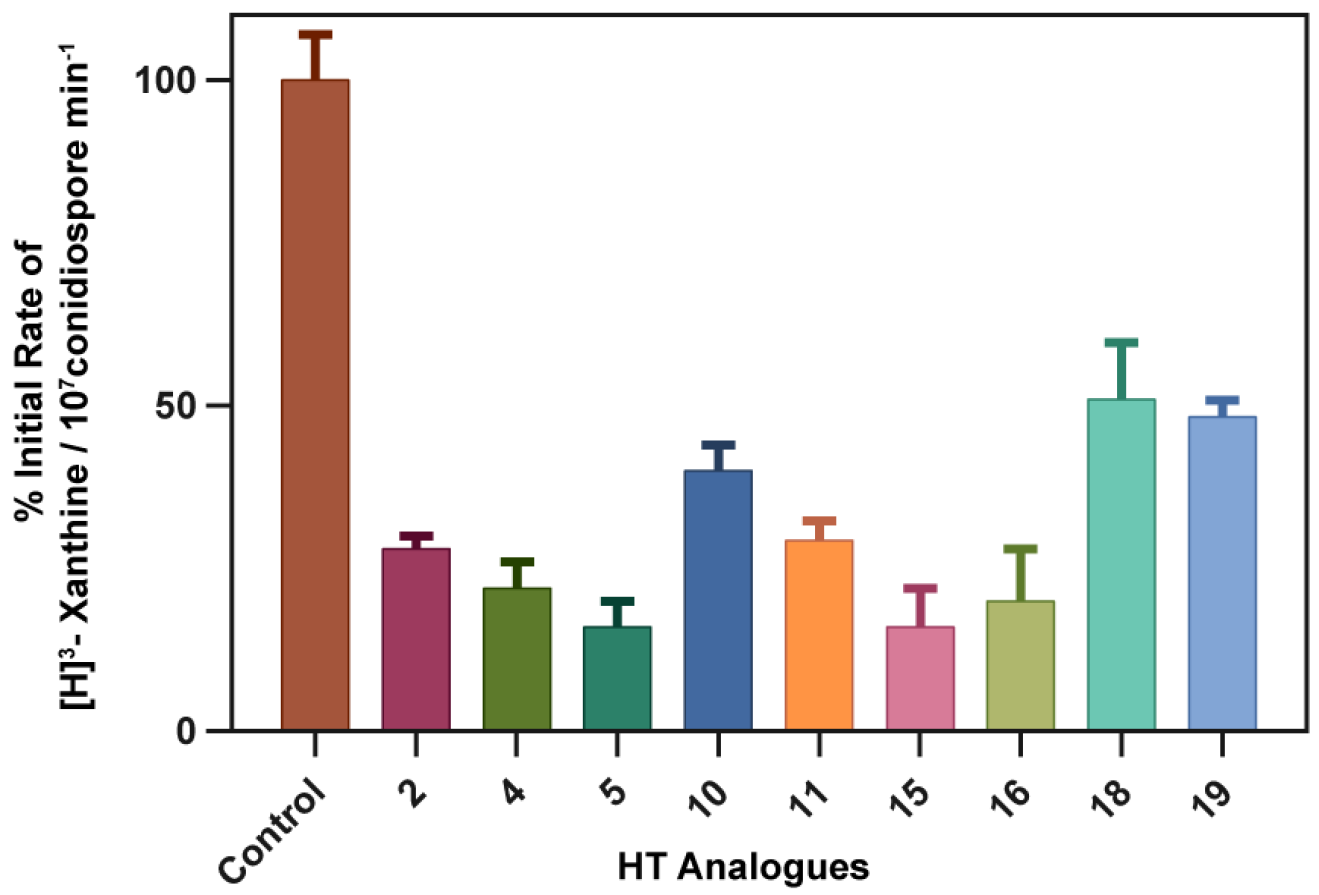
Effect of HT analogues on transporter-mediated uptake of xanthine. % of initial uptake rates are shown as recorded in a standard *A. nidulans* strain. Analogues were added at final concentration of 100 μM 10 min prior to uptake measurements. Control stands for a sample where only the solvent DMSO was added at a concentration identical to the one used as a solvent in the analogue samples. Control values were arbitrary taken as 100% rate. The results shown represent averages if three independent assays with SD <15%.

### Lack of resistance to HT analogue antifungal action

Based on our findings that showed that HT analogues act directly and rapidly on the PM, we presumed that resistance to this novel type of antifungals will be infrequent if any, similar to the case of other antimicrobials that target directly the periphery of microbes, such as amphotericin B (plasma membrane) or Echinocandins (cell wall)(Cuenca-Estrella 2014). To test this, we performed several experiments of standard u.v. or transposon-driven mutagenesis using 10^9-10^ conidiospores of an appropriate *A. nidulans* strain (i.e. one carrying the Minos transposable element; Evangelinos et al, 2015), and tried to select mutant colonies resistant to 400 μM of analogue 15. We did not manage to isolate any resistance colony. This apparently negative result is fully compatible with HT analogues targeting the PM of *A. nidulans*.

### Antifungal HT analogues do not elicit cytotoxicity in a mammalian cell line

N2A mouse neuroblastoma cells were used, in a standard MTT assay (Berridge et al, 2005), to test the whether antifungal HT analogues elicit cytotoxicity in a standard mammalian cell line. Analogues 15 or 16 were tested at two concentrations (100 or 400 μM) as described in materials and methods. No cytotoxicity of was detected at all cases (not shown).

## Discussion

Here we showed that specific chemically synthesized HT analogues possess broad and strong antifungal activity due to an immediate primary effect on the fungal plasma membrane. We do not however exclude that the HT analogues also have secondary cytotoxic targets (e.g. cell wall or enzymes), mostly in other microorganisms, which we did not test directly here. Two additional findings classify this specific set of compounds as very promising novel antifungals; lack of detectable fungal (*A. nidulans*) resistance due to mutations, and preliminary evidence for nontoxicity against mammalian cell lines. The lack of resistance is in total agreement with the herein shown destructive effect of these compounds on the fungal membrane, and resembles cases of other antifungals targeting the plasma membrane, such as amphotericin B, where only infrequent resistance is obtained due to mutations affecting the level of ergosterol and phospholipids in the membrane (Cuenca-Estrella 2014; Fairlamb et al., 2016). Overall, our results open the way for the rigorous development of a novel, efficient and maybe safer class of antifungals. Additionally, the antifungal HT analogues were also shown to also possess antimicrobial activity against specific bacteria. It remains to be shown how HT analogues have a specific action against the membrane of all fungi and several G+ bacteria.

In several previous studies, HT has been shown to have variable antibacterial activity at relatively high concentrations, in the range of 1-4 mM (Capasso et al. 1995; Bisignano et al. 1999; Medina et al. 2006; 2007; Tafesh et al. 2011; Medina-Martímez et al. 2016). However, bacterial growth was never fully inhibited and a general conclusion on the effects of HT on Gram-positive or Gram-negative species could not be made (Medina-Martínez et al. 2016). To our knowledge, the only investigations regarding the antifungal potential of HT are those against medically important *Candida* spp., which showed minimal MIC values ranging between 633 μM and 40 mM (Pereira et al., 2007; Zoric et al., 2013). Notably, fluorescent dye-exclusion based studies with *Candida albicans* revealed a membrane associated antifungal mechanism at sub-inhibitory concentrations (Zoric et al., 2013), in line with results presented herein. In the past, HT-derived compounds have been synthesized aiming at analogues with a better hydrophilic/lipophilic balance (HLB) to increase their cellular uptake and thus enhance their antioxidant or other unknown activities (Grasso et al., 2007; Bernini et al., 2012; 2015). However, no previous studies evaluating synthesized HT analogues as antifungal agents have been reported. HT derivatives have only recently been shown to have a very promising antitrypanosomal and antileishmanial activity (Belmonte-Reche et al., 2016). IC_50_ values against *Trypanosoma brucei* for HT decanoate ester and HT dodecanoate ester were 0.6 and 0.36 μM, respectively. This represented a significant 79-132 fold improvement in activity compared to HT. Focusing on structure-activity relationships, the authors found via rational design several more HT analogues to have targeted cytotoxic activity against *T. brucei* with IC_50_ values in the low micromolar range. They concluded that the di-ortho phenolic ring and medium size alkyl chain are essential for activity, whereas the nature of the chemical bond among them seems less important. Importantly, antiprotozoan HT analogues displayed a high selectivity index against MRC-5, a non-tumoral human cell line, which is in line with the targeted antimicrobial specificity we also found for the HT antifungal analogues reported here. Thus, HT is indeed a highly promising mother compound to develop novel broad range antimicrobial agents.

Several of the new compounds described here showed varying degrees of antifungal activity against the tested pathogens, nevertheless compounds **2**, **5**, **11**, **15**, **16** and **19** possessed the strongest activity, clearly higher than the mother compound HT. All the active compounds belong to the first and second series, while compounds of the third series (**21**, **22**, **23**, **24**; see **Supplementary Figure 1**) were non-active. The carbonyl substituted compounds of the first series, and especially the alkyl-substituted analogues **2**, **5** and **11**, were among the most potent compounds, suggesting that the substitution on the α-carbon is crucial for the activity. It seems that the presence of the hydroxyl group, is incompatible with activity, whereas the carbonyl group increases the antifungal capacity, probably by increasing the acidity of the catechol moiety. Notably also, all compounds in this study with an aromatic substitution (**3**, **6**, **14** and **22**; see **Supplementary Figure 1**) showed no activity against the tested pathogens, while the alkyl substituted compounds of first and second series were shown to be potent antifungals. The role of the double bond of the alkyl side chain is detrimental for the activity (see **Supplementary Figure 1**). Additionally, the role of the branching is not clear since the branched alkyl chained compounds **2**, **5**, **10**, **11**, **16**, **18** and **19** showed potent antifungal activity, whereas compounds **9**, **12**, **13** and **20** seem to be inactive. Nevertheless, our results demonstrate that only the alkyl substituted compounds of first and second series are active against the tested pathogens, with the most effective being those with an alkyl chain of 6 to 10 carbons. These results are also in agreement with the ones against Trypanosoma brucei for HT decanoate ester and HT dodecanoate ester (Belmonte-Reche et al., 2016).

Probably, the aliphatic ester side chain (hydrophobic tail) interferes with the cell membrane while the catechol group (hydrophilic head) seems to be essential for the antifungal activity. Apparently, the initial interaction involves the electrostatic interaction of the catechol system with the negatively charged phosphate groups of the fungal lipid bilayer membrane; thus the acidity of the hydroxyl groups might be essential for activity. The alkyl tail seems also crucial for antifungal activity, indicating that the interaction with the alkyl chain plays a vital role in the first step in the action of the compounds. The length (6 to 10 carbons) and the branching seem crucial for the interaction with the fungal membrane, whereas the substitution of the α-carbon is vital for the electrostatic interaction of the hydrophilic head. Apparently, these analogues, due their amphiphilic character, are inserted into the lipid bilayer of fungal and possibly other microbial membranes, where they elicit an immediate destruction effect.

Although recent technological advances made accessible unprecedented tools for drug discovery the development of new molecular scaffolds with demonstrated pharmacological properties is becoming slower and more expensive. Our study shows once more that the natural product chemical space provides the basis of successful design of new small-molecular-weight molecules with excellent pharmacological properties. As HT has been shown to undergo rapid oxidation the antifungal HT analogues described here will however need to be tested *in vivo* in mouse models and evaluated in respect to their stability or synergistic or antagonistic effects with other antifungals.

## Materials and methods

### Synthesis of HT analogues

Chemical synthesis of HT analogues is described in detail in Supplementary material. HT analogues were prepared in DMSO and aliquots and kept at −20 °C. For experiments, the final concentration of DMSO in the medium was < 0.1% (v/v) and the controls received DMSO only.

### Fungal and bacterial strains and growth media

Standard ‘wild-type’ fungal strains of *A. nidulans, A. fumigatus, A. flavus F. oxysporum* and *C. albicans* were used. The *A. nidulans* used contains a mutant allele (*veA1*) at the *velvet* locus, and a standard vitamin auxotrophy (*pabA1* for para-aminobenzoic acid requirement). All other Aspergilli and *C. albicans* used correspond to standard strains used for genome sequencing (see http://genome.jgi.doe.gov/programs/fungi/index.jsf). For *in vivo* epifluorescence microscopy, strains expressing GFP-tagged functional UapA (Pantazopoulou et al., 2007) or FurA (Krypotou et al., 2015) transporters were used. Standard Aspergillus Minimal and Complete Media (MM and CM) were used for growth of all fungi (http://www.fgsc.net). The nitrogen source used was 10 mM ammonium tartrate or NaNO_3_. Standard bacterial strains, coming from an in-house stock, *of E. coli (DH5a), P. aeruginosa, P. fluorescens, Klebsiella sp., B. subtilis* and *S. aureus* were used. Luria-Bertani medium was used for growth of bacterial strains.

### *In vivo* evaluation of HT analogues as antifungals

Fungal strains of *A. nidulans, A. fumigatus, A. flavus, F. oxysporum* and *C. albicans* were tested for their sensitivity/resistance to different concentrations. Fungal spores of each strain were used to centrally inoculate a series of 35mm petri dishes containing Minimal Media (MM) with NaNO_3_ as nitrogen source, in the presence or absence of various HT analogue concentrations dissolved in DMSO. Controls without HT analogues contained solely DMSO. The final concentration range tested for toxicity of analogues in this work was 0-1000 μM. Figures shown highlight results using concentrations of 100 or 400 μM. Growth was followed after for 4-6 days, at 25 or 37 ° C and different pH. Approximate Minimal Concentration leading to no evident fungal growth (apparent *MIC*), as well as, the concentration leading to 50% reduction in the radial diameter of the growing colonies, (*IC*_50_), after 2-4 days (depending on the fungus) were recorded for each analogue. For the non-filamentous fungus *C. albicans* we performed sensitivity tests in both liquid and solid cultures. For solid cultures testing, cells from a fresh liquid culture (O.D._600nm_ = 0.5 at 10 nm) were streaked on standard CM containing 100-400 μM of HT analogues, and incubated for 2 days at 37 °C. For liquid culture testing, HT analogues were added, at 100-400 μM, at the start of the exponential phase (O.D._600nm_ = 0.5 at 10 nm) of a *C. albicans* culture and O.D._600nm_ measurements were recorded hourly for the next 6 h, and after 24 h.

### *In vivo* evaluation of HT analogues as antibacterial agents

Standard bacterial strains *of E. coli (DH5a), P. aeruginosa, P. fluorescens, Klebsiella sp., B. subtilis* and *S. aureus* were tested for their sensitivity/resistance to different concentrations of compounds, which proved to act as antifungals. These tests were carried out either in solid or liquid LB media, in the presence or absence of 100-500 μM of HT analogues. Tests on solid media were performed by recording single colony growth after bacterial streaking. Tests in liquid media were performed by recording O.D._600nm_ values (10 nm) after addition of 100-200 μM of compounds in fresh exponentially growing bacterial cultures (O.D._600nm_ = 0.2-0.4), and comparing these values to control cultures with no analogues.

### *In vivo* epifluorescence microscopy

Samples for standard inverted epifluorescence microscopy of *A. nidulans* strains were prepared as previously described (Martzoukou et al., 2017). In brief, germlings were incubated in sterile 35mm μ-dishes, high glass bottom (*ibidi*, Germany) in 2 ml liquid MM with NaNO_3_ as nitrogen source and the necessary vitamin supplements for 20 h at 25 °C. DMSO (0.1%) or HT analogues (final concentrations tested 37.5 or 100 μM) dissolved in DMSO were added in samples under the microscope. Control samples where only DMSO was added (0.03-0.01%) were also evaluated. Images were taken before and immediately after the addition of the analogues (or DMSO) and for a period of up to 30 min. The strains used express the UapA or the FurA transporter as protein fluorescent markers specific for the plasma membrane (Pantazopoulou et al., 2007; Krypotou et al., 2015). Calcuofluor white staining of cell wall was as described in Martzoukou et al., 2017. Images were obtained with an AxioCam HR R3 camera using the Zen lite 2012 software. Contrast adjustment, area selection and color combining were made using the Zen 2012 software. Images exported as tiffs were annotated and further processed in Adobe Photoshop CS4 Extended version 11.0.2 software for brightness adjustment, rotation and alignment.

### Transport measurements

Kinetic analysis-[^3^H]-xanthine (21.6 Ci/mmol, Moravek Biochemicals, CA, USA) uptake in MM was assayed in germinating conidiospores of *A. nidulans* concentrated at 10^7^ conidiospores/100 μL, at 37 °C, pH 6.8, as previously described (Krypotou and Diallinas, 2014). Initial velocities were measured at 1 min of incubation with concentrations of 0.2-2.0 μM of [^3^H]-xanthine at the polarity maintenance stage (3-4 h, 130 rpm).

### Mutagenesis

UV mutagenesis was performed at a standard distance of 20 cm from an Osram HNS30 UV-B/C lamp. 10^9-10^ conidiospores of a standard wild-type *A. nidulans* strain or a strain possessing a dual transposition system based on the *Minos* element (Evangelinos et al., 2015) were irradiated for 4 min and subsequently plated on MM plus nitrate medium containing 400 μM of the HT analogue 15. No colonies appeared after 1 week incubation at 25 °C. The same experiment was performed twice.

### Toxicity in a mammalian cell line

N2A mouse neuroblastoma cells were grown in Dulbecco’s Modified Eagle’s Medium (DMEM) that contained 10% fetal bovine serum, 1% of penicillin and streptomycin in 96-well plates at a density of 15,000 cells/well. A standard MTT assay was used to assess cell metabolic activity in the absence (addition of solely DMSO) and presence of HT analogues (Berridge et al, 2005). The cultures were grown for 6 days at 37 °C with 5% CO_2_. Then the medium was changed to one containing 100 or 400 μM of 15 or 16 and incubated for 24 h at 37 °C with 5% CO_2_. In all cases the final concentration of DMSO was ≤ 0.1%. 20 μl of the dye MTT (2.5 mg/ml MTT in PBS) was added to each well and incubated for 4 h. The resulting formazan dye was extracted with 100 μl isopropanol/HCl (100 ml isopropanol + 833 μl HCl) and the absorbance was measured spectrophotometrically at a wavelength of 545 nm. Statistical analysis: All experiments were repeated three times. One-way ANOVA with Bonferroni’s Multiple Comparison Test was used to evaluate the statistical significance of the differences. Statistical significance was defined as p<0.05.

## Acknowledgments

We thank Dr. Joseph Meletiadis, Assist. Prof. of Microbiology in the Medical School, of University of Athens, for Amphotericin B, and Michalis Papadourakis for help in preliminary uptake experiments. This work was partly by supported by a “*Stavros Niarchos Foundation*” research grant and by “*Fondation Santé*”.

## Authors Contribution

GD performed and designed biological experiments, analyzed results and wrote the manuscript. NR performed biological experiments and made relevant figures. VT and IZ performed biological experiments. IK and ACK performed chemical experiments. EM and ALS analyzed results. IK designed chemical synthesis, analyzed results and wrote the manuscript.

## Supplementary material

### Experimental

Melting points were determined on a Büchi apparatus and are uncorrected. 1H NMR spectra and 2D spectra were recorded on a Bruker Avance III 600 or a Bruker Avance DRX 400 instrument, whereas 13C NMR spectra were recorded on a Bruker Avance III 600 or a Bruker AC 200 spectrometer in deuterated solvents and were referenced to TMS ( d scale). The signals of 1H and 13C spectra were unambiguously assigned by using 2D NMR techniques: 1H1H COSY, NOESY, HMQC, and HMBC. Mass spectra were recorded with a LTQ Orbitrap Discovery instrument, possessing an Ionmax ionization source. Flash chromatography was performed on Merck silica gel 60 (0.040e0.063 mm). Analytical thin layer chromatography (TLC) was carried out on precoated (0.25 mm) Merck silica gel F-254 plates.

### General Procedure for the synthesis of compounds 2-12

Sodium hydride (260 mg, 6.52 mmol, 60% in paraffin oil) was added at 0 °C, under argon to a solution of the appropriate acid (3.26 mmol) in dry DMF (20 mL) and the reaction mixture was stirred at room temperature for 5 min. The reaction was then cooled to 0 °C, a solution of 2-Chloro-3′,4′-dihydroxyacetophenone (0.91 g, 4.89 mmol) in DMF (2 mL) was added dropwise and the mixture was stirred at 70 °C for 4-8 h. After completion of the reaction, the volatiles were vacuum evaporated, the resulting residue was dissolved in ethyl acetate (60 mL) and washed with water (3 x 20 mL), saturated NaCl solution, dried (anhydrous Na_2_SO_4_) and evaporated to dryness. The residue was purified by column chromatography to afford the title compounds.

### 2-(3,4-dihydroxyphenyl)-2-oxoethyl adamantane-1-carboxylate (2)

The crude product was purified by column chromatography (silica gel) using a mixture of cyclohexane / ethyl acetate (4/1) as the eluent, affording the corresponding ester **2** in 73% yield.

Mp: 172-173 °C (Ethanol). ^1^H NMR (400 MHz, DMSO-*d6*) δ (ppm): 10.00 (br s, 1H, D_2_O exchang., 3’-O*H*), 9.45 (br s, 1H, D_2_O exchang., 4’-O*H*), 7.35 (d, *J* = 8.2 Hz, 1H, H-6’), 7.31 (s, 1H, H-2’), 6.84 (d, *J* = 8.2 Hz, 1H, H-5’), 5.31 (s, 2H, C*H*_2_O), 2.04-1.96 (m, 3H, C*H* _adamantyl_), 1.95-1.86 (m, 6H, C*H*_2 adamantyl_), 1.78-1.60 (m, 6H, C*H*_2 adamantyl_). ^13^C NMR (151 MHz, DMSO-*d6*) δ (ppm): 190.70 (*C*OCH_2_), 175.96 (O*C*O), 151.23 (C-4’), 145.39 (C-3’), 125.86 (C-1’), 120.98 (C-6’), 115.16 (C-5’), 114.47 (C-2’), 65.56 (*C*H_2_O), 39.99 (C _adamantyl_), 38.46 (*C*H_2 adamantyl_), 35.94 (*C*H_2 adamantyl_), 27.29 (*C*H _adamantyl_). HR-MS (ESI) m/z: Calcd for C_19_H_21_O_5_: [M1 - H]^−^ = 329.1394, found 329.1386.

### 2-(3,4-dihydroxyphenyl)-2-oxoethyl 3,5-dihydroxybenzoate (3)

The crude product was purified by column chromatography (silica gel) using a mixture of cyclohexane / ethyl acetate (1/1) as the eluent, affording the corresponding ester **3** in 64% yield

Mp: 275-276 °C (Dec.) (EtOAc). ^1^H NMR (400 MHz, DMSO-*d6*) δ (ppm): 9.76 (br s, 4H, D2O exchang., 3-O*H*, 5-O*H*, 3-O*H*’, 4’-O*H*), 7.41 (d, *J* = 8.3 Hz, 1H, H-6’), 7.36 (s, 1H, H-2’), 6.89 (s, 2H, H-2, H-6), 6.87 (d, *J* = 8.3 Hz, 1H, H-5’), 6.49 (s, 1H, H-4), 5.55 (s, 2H, C*H*_2_O). ^13^C NMR (151 MHz, DMSO-*d6*) δ (ppm): 190.75 (*C*OCH_2_), 165.40 (O*C*O), 158.64 (C-3, C-5), 151.45 (C-4’), 145.55 (C-3’), 131.07 (C-1), 125.88 (C-1’), 121.19 (C-6’), 115.34 (C-5’), 114.58 (C-2’), 107.48 (C-4), 107.38 (C-2, C-6), 66.54 (*C*H2O). HR-MS (ESI) m/z: Calcd for C_15_H_11_O_7_: [M1 - H]^−^ = 303.0510, found 303.0500.

### 2-(3,4-dihydroxyphenyl)-2-oxoethyl octanoate (4)

The crude product was purified by column chromatography (silica gel) using a mixture cyclohexane / ethyl acetate (4/1) as the eluent, affording the corresponding ester **4** in 78% yield.

Mp: 105–106 °C (Et_2_O/n-hexane). ^1^H NMR (400 MHz, CDCl_3_) δ (ppm): 7.58 (d, *J* = 2.0 Hz, 1H, H-2’), 7.41 (dd, *J* = 8.3, 2.0 Hz, 1H, H-6’), 6.93 (d, *J* = 8.3 Hz, 1H, H-5’), 5.35 (s, 2H, C*H*_2_O), 2.53 (t, *J* = 7.6 Hz, 2H, 2-C*H*_2_), 1.76-1.66 (m, 2H, 3-C*H*_2_), 1.42-1.24 (m, 8H, 4-CH_2_, 5-CH_2_, 6-CH_2_, 7-CH_2_), 0.90 (t, *J* = 7.6 Hz, 3H, CH_3_). ^13^C NMR (151 MHz, CDCl_3_) δ (ppm): 191.67 (*C*OCH2), 174.46 (O*C*OCH_2_), 150.29 (C-4’), 143.86 (C-3’), 126.94 (c-1’), 122.43 (C-6’), 114.97 (C-5’), 114.61 (C-2’), 65.83 (*C*H_2_O), 34.02 (C-2), 31.63 (C-6), 29.06 (C-5), 28.91 (C-4), 24.85 (C-3), 22.59 (C-7), 14.05 (C-8). HR-MS (ESI) m/z: Calcd for C_16_H_21_O_5_: [M1 - H]^−^ = 293.1394, found 293.1386.

### 2-(3,4-dihydroxyphenyl)-2-oxoethyl 2-((1S,3s)-adamantan-1-yl)acetate (5)

The crude product was purified by column chromatography (silica gel) using a mixture of cyclohexane / ethyl acetate (2/1) as the eluent, affording the corresponding ester **5** in 88% yield.

^1^H NMR (600 MHz, CDCl_3_) δ (ppm): 7.54 (d, *J* = 2.0 Hz, 1H, H-2’), 7.40 (dd, *J* = 8.3, 2.0 Hz, 1H, H-6’), 7.03 (br s, 1H, D_2_O exchang., 3’-O*H*), 6.90 (d, *J* = 8.3 Hz, 1H, H-5’), 6.40 (br s, 1H, D_2_O exchang., 4’-O*H*), 5.30 (s, 2H, *CH*_2_O), 2.26 (s, 2H, *CH*_2_CO), 1.98 (m, 3H, *CH* _adamantyl_), 1.74–1.59 (m, 12H, *CH*_2 adamantyl_). ^13^C NMR (151 MHz, CDCl_3_) δ (ppm): 191.77 (*C*OCH_2_), 172.39 (O*C*OCH_2_), 150.34 (C-4’), 143.98 (C-3’), 127.22 (C-1’), 122.56 (C-6’), , 115.10 (C-5’), 114.77 (C-2’), 65.81 (*C*H_2_O), 48.74 (*C*H_2_CO), 42.45 (*C*H_2 adamantyl_), 36.83 (*C*H_2 adamantyl_), 33.17 (C _adamantyl_), 28.77 (*C*H _adamantyl_). HR-MS (ESI) m/z: Calcd for C_20_H_22_O_5_: [M1 - H]^−^ = 343.1550, found 343.1546.

### (E)-2-(3,4-dihydroxyphenyl)-2-oxoethyl 3-(4-hydroxy-3-methoxyphenyl)acrylate (6)

The crude product was purified by column chromatography (silica gel) using a mixture of cyclohexane / ethyl acetate (1/5) as the eluent, affording the corresponding ester **6** in 75% yield.

^1^H NMR (600 MHz, CDCl_3_-MeOD) δ (ppm): 7.62 (d, *J* = 15.9 Hz, 1H, COCH=C*H*), 7.40 (d, *J* = 2.0 Hz, 1H, H-2’), 7.38 (dd, *J* = 8.3 Hz, 2.0 Hz, 1H, H-6’), 7.03-7.06 (m, 2H, H-2, H-6), 6.83-6.87 (m, 2H, H-5, H-5’), 6.33 (d, *J* = 15.8 Hz, 1H, COC*H*=CH), 5.30 (s, 2H, CH_2_O), 3.82 (s, 3H, OC*H*_3_). ^13^C NMR (151 MHz, CDCl_3_-MeOD) δ (ppm): 193.11 (*C*OCH_2_), 168.28 (O*C*OCH), 152.48 (C-4’), 150.29 (C-4), 148.95 (c-5), 147.42 (C-3’), 146.32 (COCH=CH), 127.37 (C-1’), 127.29 (C-1), 124.02 (C-6’), 122.44 (C-2), 116.23 (C-5’), 115.83 (COCH=CH), 115.38 (C-3), 114.44 (C-2’), 111.41 (C-6), 66.70 (*C*H_2_O), 56.32 (OCH_3_). HR-MS (ESI) m/z: Calcd for C_18_H_15_O_7_: [M1 - H]^−^ = 343.0823, found 343.0817.

### (E)-2-(3,4-dihydroxyphenyl)-2-oxoethyl docos-13-enoate (7)

The crude product was purified by column chromatography (silica gel) using a mixture of cyclohexane / ethyl acetate (4/1) as the eluent, affording the corresponding ester **7** in 76% yield.

^1^H NMR (600 MHz, CDCl_3_) δ (ppm): 7.55 (d, *J* = 2.0 Hz, 1H, H-2’), 7.40 (dd, *J* = 8.3, 2.0 Hz, 1H, H-6’), 6.91 (d, *J* = 8.3 Hz, 1H, H-5’), 6.83 (br s, 1H, D_2_O exchang., 3’-O*H*), 6.26 (br s, 1H, D_2_O exchang., 4’-O*H*), 5.38-5.32 (m, 2H, C*H*=C*H*), 5.31 (s, 2H, C*H*_2_O), 2.51 (t, *J* = 7.6 Hz, 2H, COC*H*_2_), 2.04-1.98 (m, 4H, 12-C*H*_2_ 15-C*H*_2_), 1.70 (m, 2H, 3-C*H*_2_), 1.40-1.17 (m, 28H, C*H*_2 erucic_), 0.88 (t, *J* = 7.0 Hz, 3H, C*H*_3_). ^13^C NMR (151 MHz, CDCl_3_) δ (ppm): 191.56 (*C*OCH_2_), 174.36 (O*C*OCH_2_), 150.26 (C-4’), 143.93 (C-3’), 130.06 (*C*H=*C*H), 127.25 (C-1’), 122.56 (C-6’), 115.11 (C-5’), 114.74 (C-2’), 65.92 (*C*H_2_O), 34.17 (C-2), 32.06 (C-20), 29.93 (C*H*_2erucic_), 29.86 (*C*H_2 erucic_), 29.77 (*C*H_2 erucic_), 29.76 (*C*H_2 erucic_), 29.72 (*C*H_2 erucic_), 29.68 (*C*H_2 erucic_), 29.61 (*C*H_2 erucic_), 29.47 (*C*H_2 erucic_), 29.42 (*C*H_2 erucic_), 29.28 (*C*H_2 erucic_), 27.37 (C-12, C-15), 25.02 (C-3), 22.83 (C-21), 14.26 (*C*H_3_). HR-MS (ESI) m/z: Calcd for C_30_H_47_O_5_: [M1 - H]^−^ = 487.3429, found 487.3420.

### (E)-2-(3,4-dihydroxyphenyl)-2-oxoethyl penta-2,4-dienoate (8)

The crude product was purified by column chromatography (silica gel) using a mixture of CH_2_Cl_2_ / MeOH (100/0.75) as the eluent, affording the corresponding ester **8** in 80% yield.

^1^H NMR (600 MHz, Acetone-*d*_6_) δ δ (ppm): 7.31–7.37 (m, 2H, H-2’, H-6’), 7.18 (m, 1H, COCH=C*H*), 6.83 (d, *J* = 8.3 Hz, 1H, H-5’), 6.25–6.09 (m, 2H, COC*H*=CH, CH_3_CH=C*H*), 5.82 (m, 1H, CH_3_C*H*=CH), 1.73 (t, *J* = 5.1 Hz, 1H, C*H*_3_). ^13^C NMR (151 MHz, CDCl_3_) δ (ppm): 191.28 (*C*OCH_2_), 166.70 (O*C*OCH), 151.62 (C-4’), 146.42 (*C*H _sorbic_), 146.11 (C-3’), 140.57 (*C*H _sorbic_), 130.69 (*C*H _sorbic_), 128.05 (C-1’), 122.21 (C-6’), 119.28 (*C*H _sorbic_), 115.91 (C-5’), 115.33 (C-2’), 66.49 (*C*H_2_O), 18.69 (*C*H_3_). HR-MS (ESI) m/z: Calcd for C_14_H_13_O_5_: [M1 - H]^-^ = 261.0768, found 261.0770.

### 2-(3,4-dihydroxyphenyl)-2-oxoethyl 4-methylenecyclohexanecarboxylate (9)

The crude product was purified by column chromatography (silica gel) using a mixture of CH_2_Cl_2_ / MeOH (100/1.5) as the eluent, affording the corresponding ester **9** in 85% yield.

^1^H NMR (600 MHz, CDCl_3_) δ (ppm): 7.51 (d, *J* = 2.0 Hz, 1H, H-2’), 7.40 (dd, *J* = 8.3, 2.1 Hz, 1H, H-6’), 6.91 (d, *J* = 8.3 Hz, 1H, H-5’), 6.60 (br s, 1H, D_2_O exchang., 3’-O*H*), 6.30 (br s, 1H, D_2_O exchang., 4’-O*H*), 5.30 (s, 2H, C*H*_2_O), 4.67 (s, 2H, *CH*_2_=C, 2.67 (m, 1H, COC*H*), 2.44-2.35 (m, 2H, C*H* _cyclohexyl_), 2.14-2.07 (m, 4H, C*H* _cyclohexyl_), 1.73-1.64 (m, 2H, C*H* _cyclohexyl_). ^13^C NMR (151 MHz, CDCl_3_) δ (ppm): 191.23 (*C*OCH_2_), 175.67 (O*C*OCH), 150.08 (C-4’) ,147.60 (C=*C*H_2_), 143.91 (C-3’), 127.40 (C-1’), 122.51 (C-6’),115.09 (C-5’),114.69 (C-2’), 108.24 (C=*C*H_2_), 68.16 (*C*HCO), 65.84 (*C*H_2_O), 42.53(C _cyclohexyl_), 33.70 (C _cyclohexyl_), 30.26(C _cyclohexyl_), 25.76 (C _cyclohexyl_). HR-MS (ESI) m/z: Calcd for C_16_H_17_O_5_: [M1 - H]^−^ = 289.1081, found 289.1079.

### 2-(3,4-dihydroxyphenyl)-2-oxoethyl 4-isopropylcyclohexanecarboxylate (10)

The crude product was purified by column chromatography (silica gel) using a mixture of CH_2_Cl_2_ / MeOH (100/1) as the eluent, affording the corresponding ester **10** in 78% yield.

^1^H NMR (600 MHz, CDCl_3_) δ (ppm): δ 7.38 (d, *J* = 2.0 Hz, 1H, H-2’), 7.29 (dd, *J* = 8.3, 2.0 Hz, 1H, H-6’), 6.80 (d, *J* = 8.3 Hz, 1H, H-5’), 5.29 - 5.20 (m, 2H, C*H*_2_O), 2.46 (m, 1H, CH), 2.26 (m, 1H, CH), 2.14 - 1.98 (m, 1H, CH), 1.23 (m, 1H, Ch), 1.11 (m, 1H, CH), 1.04-0.98 (m, 3H, CH_3_), 0.87 (d, *J* = 1.9 Hz, 9H, CH_3_). ^13^C NMR (151 MHz, CDCl_3_) δ (ppm): δ 192.06 (*C*OCH_2_), 173.51 (O*C*OCH_2_), 150.69 (C-4’), 144.53 (C-3’), 126.70 (C-1’), 122.13 (c-6’), 115.03 (C-5’), 114.60 (C-2’), 65.75 (*C*H_2_O), 50.61 (*C*H_2_CH), 43.58 (*C*H_2_CO), 31.06 (C), 29.96 (3xC*H*_3_), 27.03 (*C*H), 22.60 (*C*H_3_). HR-MS (ESI) m/z: Calcd for C_18_H_23_O_5_: [M1 - H]^−^ = 319.1551, found 319.1542.

### 2-(3,4-dihydroxyphenyl)-2-oxoethyl 3,5,5-trimethylhexanoate (11)

The crude product was purified by column chromatography (silica gel) using a mixture of cyclohexane / CH_2_Cl_2_ (1/10) as the eluent, affording the corresponding ester **11** in 89% yield.

^1^H NMR (600 MHz, CDCl3) δ (ppm): 7.38 (d, J = 2.0 Hz, 1H, H-2’), 7.29 (dd, *J* = 8.3, 2.0 Hz, 1H, H-6’), 6.80 (d, *J* = 8.3 Hz, 1H, H-5’), 5.29–5.20 (m, 2H, C*H*_2_O), 2.46 (m, 1H, C*H*), 2.26 (m, 1H, *C*H), 2.14–1.98 (m, 1H, C*H*), 1.23 (m, 1H, C*H*), 1.11 (m, 1H, C*H*), 1.04–0.98 (m, 3H, C*H*_3_), 0.87 (d, J = 1.9 Hz, 9H, C*H*_3_). ^13^C NMR (151 MHz, CDCl_3_) δ (ppm): 192.06 (*C*OCH_2_), 173.51 (O*C*OCH_2_), 150.69 (C-4’), 144.53 (C-3’), 126.70 (C-1’), 122.13 (C-6’), 115.03 (C-5’), 114.60 (C-2’), 65.75 (*C*H_2_O), 50.61 (C-4), 43.58 (C-2), 31.06 (C-5), 29.96 (*C*H_3_), 26.99 (C-3), 22.60 (*C*H_3_). HR-MS (ESI) m/z: Calcd for C_17_H_23_O_5_: [M1 - H]^−^ = 307.1550, found 307.1540.

### 2-(3,4-dihydroxyphenyl)-2-oxoethyl 2-cyclohexylacetate (12)

The crude product was purified by column chromatography (silica gel) using a mixture of cyclohexane / ethyl acetate (4/1) as the eluent, affording the corresponding ester **12** in 95% yield, as an oil.

^1^H NMR (400 MHz, CDCl_3_) δ (ppm): 6.81 (d, *J* = 8.0 Hz, 1H, H-5’), 6.75 (d, *J* = 1.8 Hz, 1H, H-2’), 6.62 (dd, *J* = 8.0, 1.8 Hz, 1H, H-6’), 4.27 (t, *J* = 7.0 Hz, 2H, CH_2_C*H*_2_O), 2.84 (t, *J* = 7.0 Hz, 2H, C*H*_2_CH_2_O), 2.19 (d, *J* = 7.1 Hz, 2H, COC*H*_2_), 1.82-1.60 (m, 6H, H-1, H-2, H-3, H-4, H-5, H-6), 1.31-1.08 (m, 3H, H-3, H-4, H-5), 0.98-0.91 (m, 2H, H-2, H-6). ^13^C NMR (151 MHz, CDCl_3_) δ (ppm): 174.33 (*C*O), 143.89 (C-3’), 142.58 (C-4’), 130.18 (C-1’), 121.07 (C-6’), 115.83 (C-5’), 115.31 (C-2’), 65.23 (CH_2_*C*H_2_O), 42.29 (CO*C*H_2_), 34.89 (*C*H_2_CH_2_O), 34.41 (C-1), 32.94 (C-2, C-6), 26.90 (C-4), 26.06 (C-3, C-5). HR-MS (ESI) m/z: Calcd for C_16_H_19_O_5_: [M1 - H]^−^ = 291.1237, found 291.1237.

## General Procedure for the synthesis of compounds 13-20

Triethylsilane (0.312 mL, 1.96 mmol) was added dropwise to a suspension of the appropriate ester **2-12** (0.49 mmol) in trifluoroacetic acid (0.190 mL, 2.45 mmol), at 0 °C. The flask was sealed and the resulting mixture was stirred at room temperature for 3-6 h. After completion of the reaction, the volatiles were vacuum evaporated, the resulting residue was dissolved in ethyl acetate (40 mL) and washed with water (3 × 15 mL), saturated NaCl solution, dried (anhydrous Na_2_SO_4_) and evaporated to dryness.

### 3.4-dihydroxyphenethyl adamantane-1-carboxylate (13)

The crude product was purified by column chromatography (silica gel) using a mixture of cyclohexane / ethyl acetate (4/1) as the eluent, affording the corresponding ester **13** in 77% yield.

Mp: 151-152 °C (c-Hex). ^1^H NMR (400 MHz, DMSO-*d6*) δ (ppm): 8.78 (br s, 1H, D_2_O exchang, 4’-O*H*), 8.69 (br s, 1H, D_2_O exchang, 3’-O*H*), 6.64 (d, *J* = 7.9 Hz, 1H, H-5’), 6.61 (d, *J* = 1.5 Hz, 1H, H-2’), 6.46 (dd, *J* = 7.9, 1.5 Hz, 1H, H-6’), 4.10 (t, *J* = 6.8 Hz, 2H, CH_2_C*H*_2_O), 2.68 (t, *J* = 6.8 Hz, 2H, C*H*_2_CH_2_O), 2.01-1.90 (m, 3H, *CH* _adamantyl_), 1.81-1.73 (m, 6H, C*H*_2 adamantyl_), 171-1.60 (m, 6H, C*H*_2 adamantyl_). ^13^°C NMR (50 MHz, DMSO-*d6*) δ (ppm): 176.83 (CO), 145.48 (C-3’), 143.92 (C-4’), 129.12 (C-1’), 119.98 (C-6’), 116.72 (C-2’), 115.88 (C-5’), 64.83 (CH_2_*C*H_2_O), 40.00 (C _adamantyl_), 38.82 (*C*H_2 adamantyl_), 36.39 (*C*H_2 adamantyl_), 34.32 (*C*H_2_CH_2_O), 27.76 (*C*H _adamantyl_). HR-MS (ESI) m/z: Calcd for C_19_H_23_O_4_: [M1 - H]^−^ = 315.1601, found 315.1599.

### 3,4-dihydroxyphenethyl 3,5-dihydroxybenzoate (14)

The crude product was purified by column chromatography (silica gel) using a mixture of cyclohexane / ethyl acetate (1/1) as the eluent, affording the corresponding ester **14** in 84% yield.

Mp: 110-111 °C (EtOAc-c-Hex). ^1^H NMR (600 MHz, DMSO-*d6*) δ (ppm): 9.64 (br s, 2H, D_2_O exchang, 3-O*H* ,5-O*H*), 8.79 (br s, 1H, D_2_O exchang, 4’-O*H*), 8.75 (br s, 1H, D_2_O exchang, 3’-O*H*), 6.80 (d, *J* = 2.1 Hz, 2H, H-2, H-6), 6.67 (d, *J* = 7.9 Hz, 1H, H-5’), 6.64 (d, *J* = 2.1 Hz, 1H, H-2’), 6.53 (dd, *J* = 7.9, Hz, 2.1 Hz, 1H, H-6’), 6.44 (t, *J* = 2.3 Hz, 1H, H-4), 4.32 (t, *J* = 6.8 Hz, 2H, CH_2_C*H*_2_O), 2.81 (t, *J* = 6.8 Hz, 2H, C*H*_2_CH_2_O). ^13^C NMR (151 MHz, DMSO-*d6*) δ (ppm): 166.28 (CO), 158.82 (C-3, C-5), 145.65 (C-3’), 144.32 (C-4’), 132.03 (C-1), 129.25 (C-1’), 119.84 (C-6’), 116.78 (C-2’), 116.11 (C-5’), 107.55 (C-2, C-4, C-6), 66.04 (CH_2_*C*H_2_O), 34.37 (*C*H_2_CH_2_O). HR-MS (ESI) m/z: Calcd for C_15_H_13_O_6_: [M1 - H]^−^ = 289.0717, found 289.0713.

### 3,4-dihydroxyphenethyl octanoate (15)

The crude product was purified by column chromatography (silica gel) using a mixture of cyclohexane / ethyl acetate (4/1) as the eluent, affording the corresponding ester **15** in 91% yield, as an oil.

^1^H NMR (400 MHz, CDCl_3_) δ (ppm): 6.81 (d, *J* = 8.0 Hz, 1H, H-5’), 6.75 (d, *J* = 1.6 Hz, 1H, H-2’), 6.62 (dd, *J* = 8.0, 1.6 Hz, 1H, H-6’), 4.26 (t, *J* = 7.2 Hz, 2H, CH_2_*CH*_2_O), 2.82 (t, *J* = 7.2 Hz, 2H, C*H*_2_CH_2_O), 2.32 (t, *J* = 7.2 Hz, 2H, 2-*CH*_2_), 1.66-1.57 (m, 2H, 3-C*H*_2_), 1.22-1.36 (m, 8H, 4-C*H*_2_, 5-C*H*_2_, 6-C*H*_2_, 7-C*H*_2_), 0.90 (t, *J* = 7.2 Hz, 3H, C*H*_3_). ^13^C NMR (151 MHz, CDCl_3_) δ (ppm): 175.01 (*C*O), 143.80 (C-3’), 142.50 (C-4’), 130.31 (C-1’), 121.19 (C-6’), 115.86 (C-2’), 115.36 (c-5’), 65.28 (CH_2_*C*H_2_O), 34.47 (*C*H_2_CH_2_O), 34.42 (C-2), 31.63 (C-6), 29.05 (C-5), 28.89 (C-4), 24.94 (C-3), 22.59 (C-7), 14.05 (C-8). HR-MS (ESI) m/z: Calcd for C_16_H_23_O_4_: [M1 - H]^-^ = 279.1601, found 279.1590.

### 3,4-dihydroxyphenethyl 2-(adamantan-1-yl)acetate (16)

The crude product was purified by column chromatography (silica gel) using a mixture of cyclohexane / ethyl acetate (4/1) as the eluent, affording the corresponding ester **16** in 95% yield.

^1^H NMR (600 MHz, CDCl_3_) δ (ppm): 6.79 (d, *J* = 8.1 Hz, 1H, H-5’), 6.75 (d, *J* = 2.0 Hz, 1H, H-2’), 6.63 (dd, *J* = 8.1, 2.0 Hz, 1H, H-6’), 4.24 (t, *J* = 7.1 Hz, 2H, CH_2_*CH*_2O_), 2.82 (t, *J* = 7.1 Hz, 2H, C*H*_2_CH_2_O), 2.06 (s, 2H, C*H*_*2*_CO), 1.93 (m, 3H, C*H* _adamantyl_), 1.70-1.57 (m, 6H, C*H*_2 adamantyl_), 1.54 (m, 6H, C*H*_2 adamantyl_). ^13^C NMR (151 MHz, CDCl3) δ (ppm): 172.66 (*C*O), 143.72 (C-3’), 142.41 (C-4’), 130.51 (C-1’), 121.25 (C-6’), 115.85 (C-2’), 115.30 (C-5’), 64.88 (CH_2_*C*H_2_O), 49.14 (*C*H_2_CO), 42.39 (*C*H_2 adamantyl_), 36.68 (*C*H_2 adamantyl_), 34.47 (*C*H_2_CH_2_O), 32.82 (C _adamantyl_), 28.61 (*C*H _adamantyl_). HR-MS (ESI) m/z: Calcd for C_20_H_25_O_4_: [M1 - H]^−^ = 329.1758, found 329.1749.

### (Z)-3,4-dihydroxyphenethyl docos-13-enoate (17)

The crude product was purified by column chromatography (silica gel) using a mixture of cyclohexane / ethyl acetate (4/1) as the eluent, affording the corresponding ester **17** in 95% yield.

^1^H NMR (600 MHz, CDCl_3_) δ (ppm): 6.78 (d, *J* = 8.0 Hz, 1H, H-5’), 6.73 (d, *J* = 2.0 Hz, 1H, H-2’), 6.63 (dd, *J* = 8.0, 2.0 Hz, 1H, H-6’), 5.35 (m,2H, C*H*=C*H*), 4.24 (t, *J* = 7.1 Hz, 2H, CH_2_C*H*_2_O), 2.81 (t, *J* = 7.1 Hz, 2H, C*H*_2_CH_2_O), 2.29 (t, *J* = 7.6 Hz, 2H, COC*H*_2_), 2.01 (m, 4H, 12-CH2, 15-C*H*_2_), 1.61 (m, 2H, 3-C*H*_2_), 1.38-1.17 (m, 28H, C*H*_2 erucic_), 0.88 (t, J = 6.9 Hz, 3H, C*H*_3_). ^13^C NMR (151 MHz, CDCl3) δ (ppm): 174.58 (*C*O), 143.81 (C-3’), 142.46 (C-4’), 130.74 (C-1’), 130.06 (*C*H=*C*H), 121.42 (C-6’), 116.02 (C-2’), 115.49 (C-5’), 65.18 (CH_2_*C*H_2_O), 34.58 (*C*H_2_CH_2_O), 33.82 (C-2), 32.00 (C-20), 30.24-28.81 (*C*H_2 erucic_), 27.37 (C-12, C-15), 25.02 (C-3), 22.81 (c-21), 14.24 (*C*H_3_). HR-MS (ESI) m/z: Calcd for C_30_H_49_O_4_: [M1 - H]^−^ = 473.3636, found 473.3636.

### 3,4-dihydroxyphenethyl 4-isopropylcyclohexanecarboxylate (18)

The crude product was purified by column chromatography (silica gel) using a mixture of CH_2_Cl_2_ / MeOH (100/1) as the eluent, affording the corresponding ester **18** in 96% yield.

^1^H NMR (600 MHz, CDCl_3_) δ (ppm): 6.78 (d, *J* = 8.1 Hz, 1H, H-5’), 6.73 (d, *J* = 2.0 Hz, 1H, H-2’), 6.63 (dd, *J* = 8.1, 2.0 Hz, 1H, H-6’), 5.60 (br s, 1H, D2O exchang., 4’-O*H*), 5.40 (br s, 1H, D_2_O exchang., 3’-O*H*), 4.22 (t, *J* = 7.1 Hz, 2H, CH_2_C*H*_2_O), 2.81 (t, *J* = 7.1 Hz, 2H, C*H*_2_CH_2_O), 2.25 - 2.14 (m, 1H, C*H*), 1.98 - 1.93 (m, 2H, C*H*), 1.80 - 1.73 (m, 2H, C*H*), 1.44 - 1.32 (m, 3H, C*H*), 1.06 - 0.93 (m, 3H, C*H*), 0.85 (d, *J* = 6.8 Hz, 6H, C*H*_3_). ^13^C NMR (151 MHz, CDCl_3_) δ (ppm): 176.82 (*C*O), 143.77 (C-3’), 142.39 (C-4’), 130.90 (C-1’), 121.47 (C-6’), 116.05 (C-2’), 115.46 (C-5’), 65.02 (CH2*C*H2O), 43.80 (*C*H), 43.42 (*C*H), 34.64 (*C*H_2_CH_2_O), 32.91 (*C*H), 29.35 (*C*H_2_), 29.01 (*C*H_2_), 19.88 (*C*H_3_). HR-MS (ESI) m/z: Calcd for C_18_H_25_O_4_: [M1 - H]^−^ = 305.1758, found 305.1749.

### 3,4-dihydroxyphenethyl 3,5,5-trimethylhexanoate (19)

The crude product was purified by column chromatography (silica gel) using a mixture of cyclohexane / CH_2_Cl_2_ (1/2) as the eluent, affording the corresponding ester **19** in 95% yield.

^1^H NMR (600 MHz, CDCl_3_) δ (ppm): 6.78 (d, *J* = 8.0 Hz, 1H, H-5’), 6.73 (d, *J* = 2.0 Hz, 1H, H-2’), 6.61 (dd, *J* = 8.0, 2.0 Hz, 1H, H-6’), 6.22 (br s, 1H, D_2_O exchang., 4’-OH), 6.13 (br s, 1H, D_2_O exchang., 3’-OH), 4.22 (t, *J* = 7.1 Hz, 2H, CH_2_C*H*_2_O), 2.80 (t, *J* = 7.1 Hz, 2H, C*H*_2_CH_2_O), 2.29 (m, 1H, C*H*), 2.10 (m, 1H, C*H*), 2.03 – 1.95 (m, 1H, C*H*), 1.20 (m, 1H, C*H*), 1.09 (m, 1H, C*H*), 0.93 (d, *J* = 6.6 Hz, 3H, CH_3_), 0.88 (s, 9H, C*H*_3_). ^13^C NMR (151 MHz, CDCl_3_) δ (ppm): 173.52 (*C*O), 143.78 (C-3’), 142.37 (C-4’), 130.95 (C-1’), 121.47 (C-6’), 116.08 (C-2’), 115.50 (C-5’), 64.99 (CH_2_*C*H_2_O), 50.68 (*C*H_2_CH), 44.21 (*C*H_2_CH), 34.65 (*C*H_2_CH_2_O), 31.19 (C), 30.07 (3 × C*H*_3_), 27.17 (*C*H), 22.78 (C*H*_3_). HR-MS (ESI) m/z: Calcd for C_17_H_25_O_4_: [M1 - H]^−^ = 293.1758, found 293.1750.

### 3,4-dihydroxyphenethyl 2-cyclohexylacetate (20)

The crude product was purified by column chromatography (silica gel) using a mixture of cyclohexane / ethyl acetate (4/1) as the eluent, affording the corresponding ester **20** in 95% yield, as an oil.

^1^H NMR (400 MHz, CDCl_3_) δ (ppm): 6.81 (d, *J* = 8.0 Hz, 1H, H-5’), 6.75 (d, *J* = 1.8 Hz, 1H, H-2’), 6.62 (dd, *J* = 8.0, 1.8 Hz, 1H, H-6’), 4.27 (t, *J* = 7.0 Hz, 2H, CH_2_C*H*_2_O), 2.84 (t, *J* = 7.0 Hz, 2H, C*H*_2_CH_2_O), 2.19 (d, *J* = 7.1 Hz, 2H, COC*H*_2_), 1.82-1.60 (m, 6H, H-1, H-2, H-3, H-4, H-5, H-6), 1.31-1.08 (m, 3H, H-3, H-4, H-5), 0.98-0.91 (m, 2H, H-2, H-6). ^13^C NMR (151 MHz, CDCl_3_) δ (ppm): 174.33 (*C*O), 143.89 (C-3’), 142.58 (C-4’), 130.18 (C-1’), 121.07 (C-6’), 115.83 (C-2’), 115.31 (C-5’), 65.23(CH_2_*C*H_2_O), 42.29 (CO*C*H_2_), 34.89 (*C*H_2_CH_2_O), 34.41 (C-1), 32.94 (C-2, C-6), 26.90 (C-4), 26.06 (C-3, C-5). HR-MS (ESI) m/z: Calcd for C_16_H_21_O_4_: [M1 - H]^−^ = 277.1445, found 277.1436.

## General Procedure for the synthesis of compounds 21-24

A solution of the appropriate ester (**2**, **3**, **4** or **11**) (1 mmol) in t-butanol (20 ml) was hydrogenated in the presence of 10% Pd/C (50 mg), under a pressure of 50 psi at room temperature for 3-4 h. After completion of the reaction, the resulting mixture was filtered through a celite pad and the filtrate was evaporated to dryness to afford the title compounds **21-24**.

### 2-(3,4-dihydroxyphenyl)-2-hydroxyethyl adamantane-1-carboxylate (21)

The crude product was purified by column chromatography (silica gel) using a mixture of cyclohexane / ethyl acetate (2/1) as the eluent, affording the corresponding ester **21** in 81% yield.

Mp: 148-149 °C (CHCl_3_/*n*-pentane). ^1^H NMR (400 MHz, DMSO-*d6*) δ (ppm): 8.80 (br s, 1H, D_2_O exchang., 3’-O*H*), 8.73 (br s, 1H, D_2_O exchang., 4’-O*H*), 6.76 (d, *J* = 1.9 Hz, 1H, H-2’), 6.67 (d, *J* = 8.0 Hz, 1H, H-5’), 6.59 (dd, *J* = 8.0, 1.9 Hz, 1H, H-6’), 5.30 (d, *J* = 4.4 Hz, 1H, CHO*H*), 4.58-4.53 (m, 1H, C*H*CH_2_), 4.00-3.90 (m, 2H, CHC*H*_2_), 1.98-1.94 (m, 3H, C*H* _adamantyl_), 1.81-1.75 (m, 6H, C*H*_2 adamantyl_), 1.71-1.61 (m, 6H, C*H*_2 adamantyl_). ^13^C NMR (151 MHz, DMSO-*d6*) δ (ppm): 176.32 (*C*O), 144.87 (c-3’), 144.42 (C-4’), 133.14 (C-1’), 117.15 (C-6’), 115.02 (C-2’), 113.74 (C-5’), (HO*C*HCH_2_), 68.41 (HOCH*C*H_2_), 40.05 (C _adamantyl_), 38.30 (*C*H_2 adamantyl_), 35.93 (*C*H_2 adamantyl_), 27.28 (*C*H _adamantyl_). HR-MS (ESI) m/z: Calcd for C_19_H_23_O_5_: [M1 - H]^−^ = 331.1551, found 331.1545.

### 2-(3,4-dihydroxyphenyl)-2-hydroxyethyl 3,5-dihydroxybenzoate (22)

The crude product was purified by column chromatography (silica gel) using a mixture of cyclohexane / ethyl acetate (1/1) as the eluent, affording the corresponding ester **22** in 76% yield.

Mp: 181-182 °C (CH_2_Cl_2_). ^1^H NMR (400 MHz, DMSO-*d6*) δ (ppm): 9.64 (br s, 2H, D_2_O exchang., 3-O*H*, 5-O*H*), 8.87 (br s, 1H, D_2_O exchang., 3’-O*H*), 8.82 (br s, 1H, D_2_O exchang., 4’-O*H*), 6.83 (s, 2H, H-2, H-6), 6.81 (s, 1H, H-2’), 6.69 (d, *J* = 8.0 Hz, 1H, H-5’), 6.65 (d, *J* = 8.0 Hz, 1H, H-6’), 6.44 (s, 1H, H-4), 5.47 (m, 1H, CHO*H*), 4.75-4.65 (m, 1H, C*H*CH_2_), 4.20-4.10 (m, 2H, CHC*H*_2_). ^13^C NMR (151 MHz, DMSO-*d6*) δ (ppm): 166.00 (*C*O), 158.46 (C-3,5), 145.00 (C-3’), 144.67 (C-4’), 132.82 (C-1’), 131.30 (C-1), 116.99 (C-6’), 115.04 (C-2’), 113.41 (C-5’), 107.26 (C-2, C-4, C-6), 69.93 (*C*HCH_2_), 69.23 (CH*C*H_2_). HR-MS (ESI) m/z: Calcd for C_15_H_13_O_7_: [M1 - H]^−^ = 305.0667, found 305.0654.

### 2-(3,4-dihydroxyphenyl)-2-hydroxyethyl octanoate (23)

The crude product was purified by column chromatography (silica gel) using a mixture of cyclohexane / ethyl acetate (2/1) as the eluent, affording the corresponding ester **23** in 87% yield.

Mp: 119-120 °C (CH_2_Cl_2_/*n*-pentane). ^1^H NMR (400 MHz, DMSO-*d6*) δ (ppm): 8.84 (br s, 1H, D_2_O exchang., 3’-O*H*), 8.77 (br s, 1H, D_2_O exchang., 4’-O*H*), 6.75 (d, *J* = 2.0 Hz, 1H, H-2’), 6.66 (d, *J* = 8.0 Hz, 1H, H-5’), 6.57 (dd, *J* = 8.0 Hz, 2.0 Hz, 1H, H-6’), 5.34 (m, 1H, CHO*H*), 4.61-4.52 (m, 1H, C*H*CH_2_), 4.00-3.90 (m, 2H, CHC*H*_2_), 2.26 (t, *J* = 7.3 Hz, 2H, 2-C*H*_2_), 1.56-1.44 (m, 2H, 3-C*H*_2_), 1.34-1.14 (m, 8H, 4-C*H*_2_, 5-C*H*_2_, 6-C*H*_2_, 7-C*H*_2_), 0.85 (t, *J* = 7.3 Hz, 3H, C*H*_3_). ^13^C NMR (151 MHz, DMSO-*d6*) δ (ppm): 172.82 (*C*O), 144.93 (C-3’), 144.51 (C-4’), 132.97 (C-1’), 117.09 (C-6’), 115.10 (C-2’), 113.68 (C-5’), 69.83 (HO*C*HCH_2_), 68.79 (HOCH*C*H_2_), 33.37 (C-2), 31.09 (C-6), 28.36 (C-4, C-5), 24.41 (C-3), 22.03 (C-7), 13.93 (C-8). HR-MS (ESI) m/z: Calcd for C_16_H_23_O_5_: [M1 - H]^−^ = 295.1550, found 295.1543.

### 2-(3,4-dihydroxyphenyl)-2-hydroxyethyl 3,5,5-trimethylhexanoate (24)

The crude product was purified by column chromatography (silica gel) using a mixture of CH_2_Cl_2_ / MeOH (100/1) as the eluent, affording the corresponding ester **24** in 94% yield, as an oil.

^1^H NMR (600 MHz, Acetone-*d6*) δ (ppm): 7.08 (br s, 1H, D_2_O exchang., O*H*), 6.92 (d, *J* = 1.9 Hz, 1H, H-2’), 6.78 (d, *J* = 8.0 Hz, 1H, H-5’), 6.74 (dd, *J* = 8.0, 1.9 Hz, 1H, H-6’), 4.76 (m, 1H, CHO*H*), 4.15-4.03 (m, 2H, C*H*CH_2_), 2.29 (m, 1H, C*H*), 2.15-2.07 (m, 2H, 2 × C*H*), 1.28 (m, 1H, C*H*), 1.10 (m, 1H, C*H*), 0.95 (d, *J* = 6.6 Hz, 3H, CH_3_), 0.91 (s, 9H, C*H*_3_). ^13^C NMR (151 MHz, Acetone-*d6*) δ (ppm): 172.11 (*C*O), 144.85 (C-3’), 144.47 (C-4’), 133.64 (C-1’), 117.75 (C-6’), 114.85 (C-2’), 113.39 (C-5’), 71.12 (HO*C*HCH_2_), 68.91 (HOCH*C*H_2_), 50.27 (*C*H_2_CH), 43.47 (*C*H_2_CH), 30.62 (C), 29.34 (3 × C*H*_3_), 26.75 (*C*H), 22.07 (*C*H_3_). HR-MS (ESI) m/z: Calcd for C_17_H_25_O_5_: [M1 - H]^−^ = 309.1707, found 309.1696.

## Supplementary Figures

**Supplementary Figure 1.**
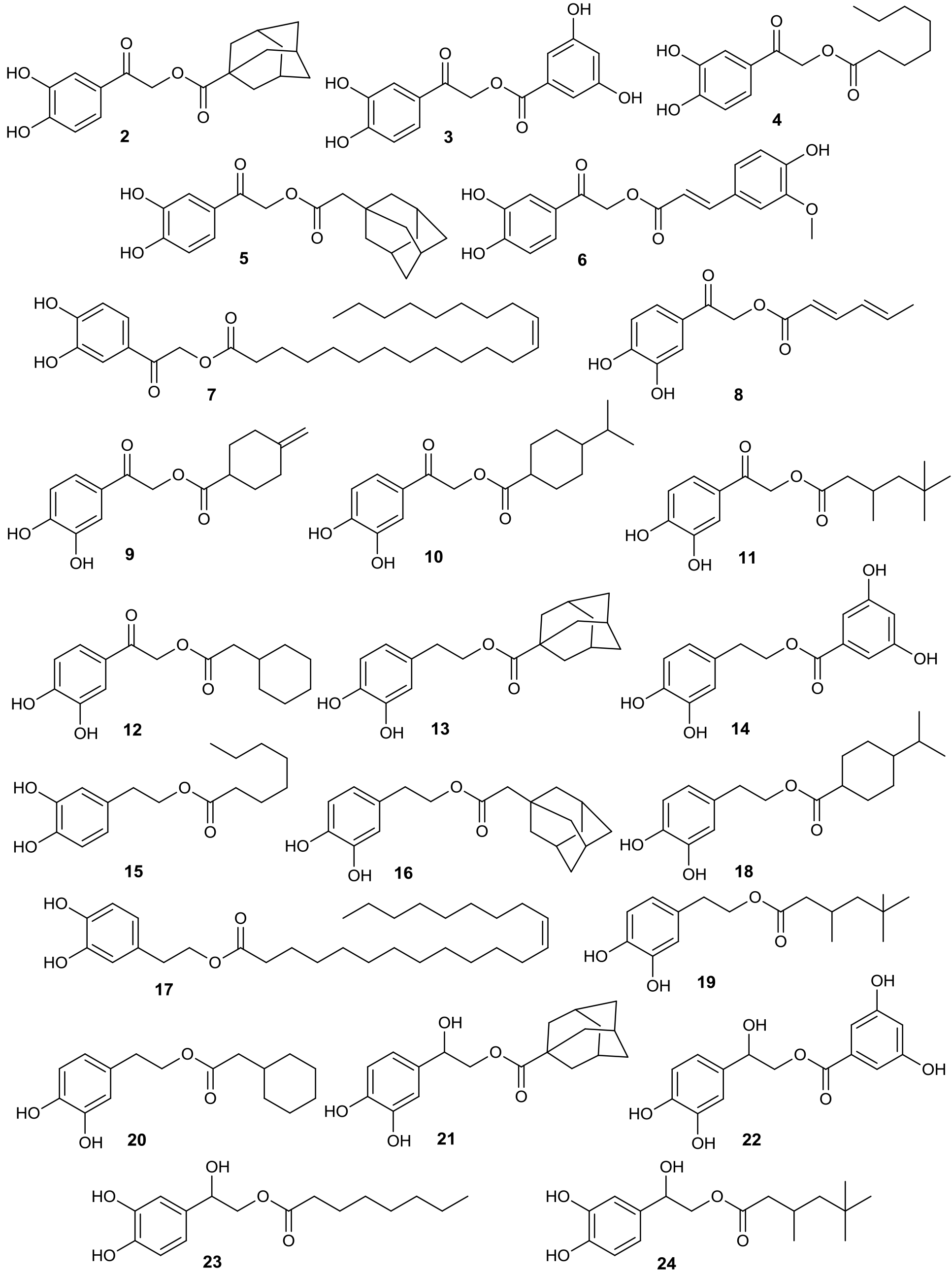
Synthesized analogues of HT

**Supplementary Figure 2.**
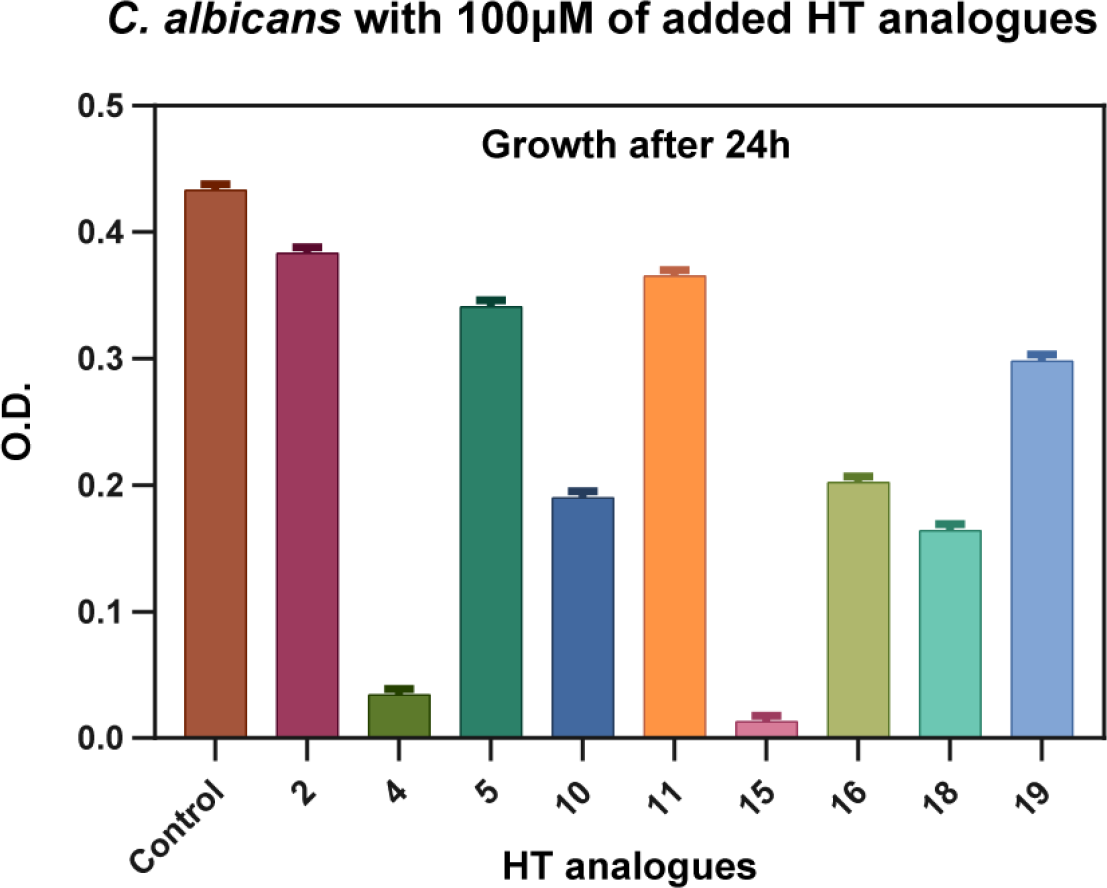
Overnight growth of *C. albicans* in the presence of HT analogues. Control stands for a culture where instead of an HT analogue, DMSO solvent was added in a concentration identical to that in which HT analogues were dissolved. Growth is recorded after 24 h as O.D. values at 600 nm.

**Supplementary Figure 3.**
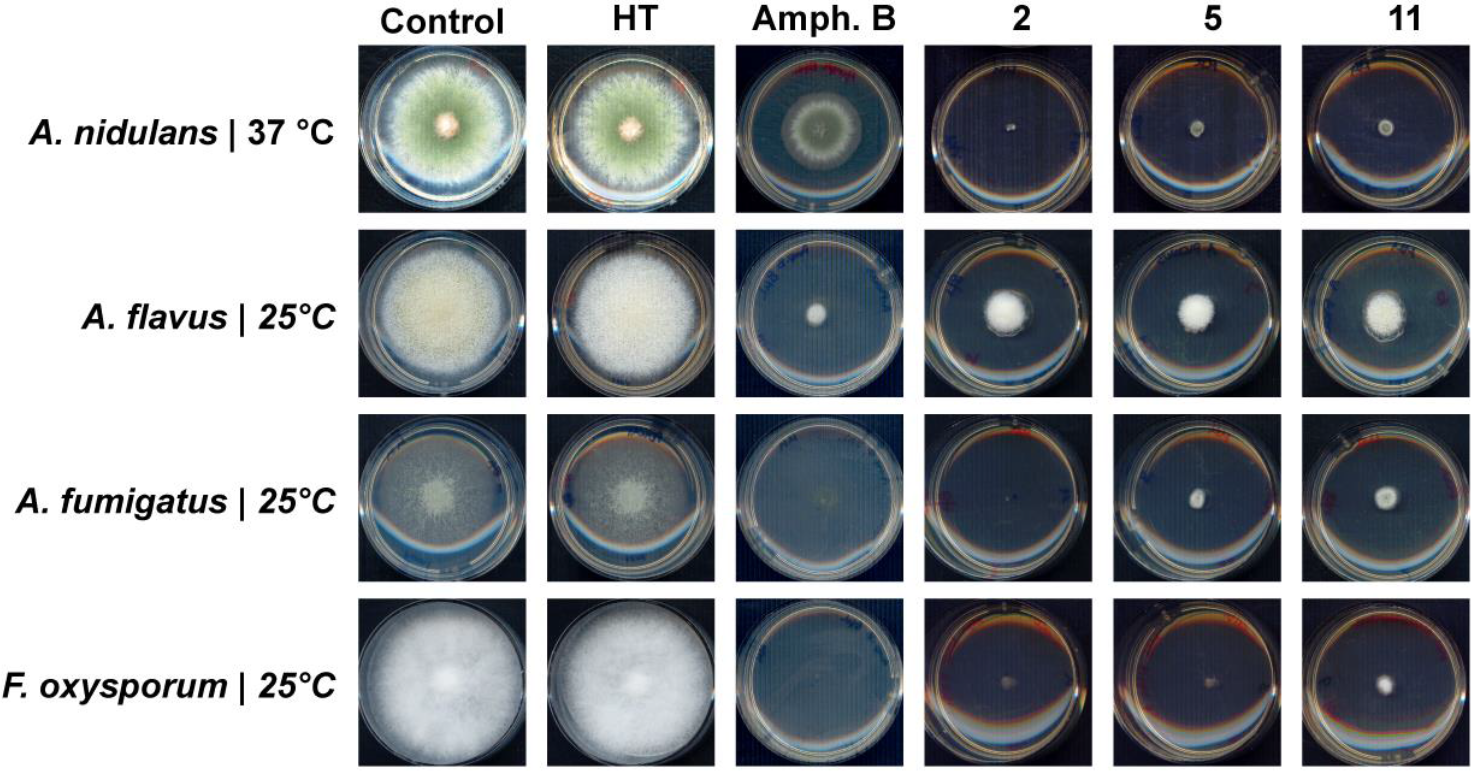
Comparison of antifungal activity of HT, amphotericin B and selected HT analogues. The concentration of antifungals shown is 100 μM. Control stands for a culture where DMSO solvent was added in a concentration identical to that of antifungals.

**Supplementary Figure 4.**
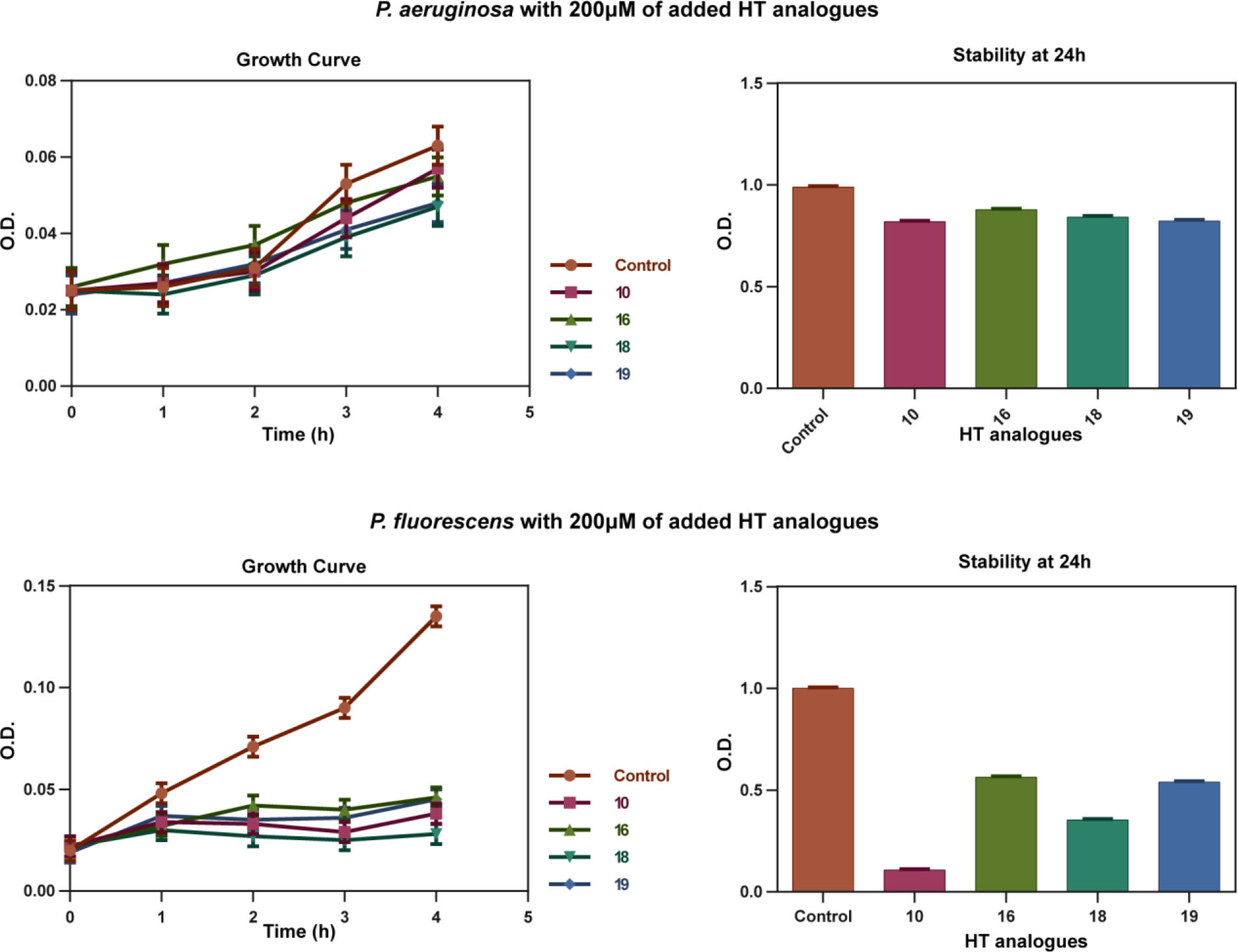
Antibacterial activity of HT analogues against *P. aeruginosa* and *P. fluorescens*. Growth curves on the left show O.D. values recorded hourly at 600 nm. Column bar graphs on the right show the growth after 24 hours after the HT analogues addition, at 600 nm. Control stands for a culture DMSO solvent was added in a concentration identical to that of HT analogues.

**Supplementary Figure 5.**
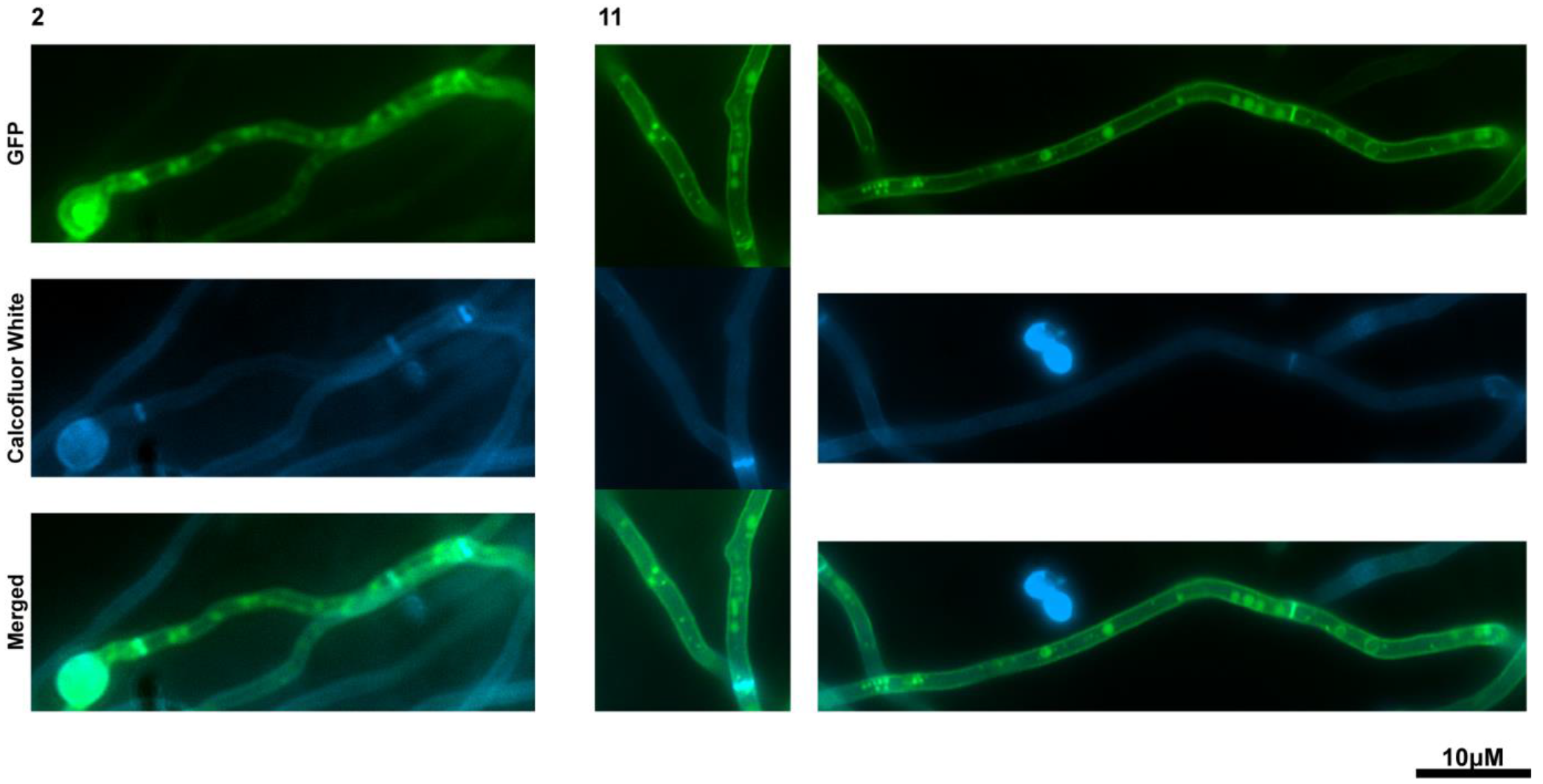
Epifluorescence *in vivo* microscopy showing the apparent non-effect of HT analogues 2 and 11 (37.5 μM) on *A. nidulans* cell wall. The picture shows hyphae of strains expressing functional, GFP-tagged, FurA as PM molecular marker, stained with Calcuofluor white as a standard marker for cell wall integrity (Martzoukou et al., 2017).

